# Crop-specific and microbially-mediated impacts of alternative agricultural amendments on plant performance

**DOI:** 10.64898/2025.11.28.691263

**Authors:** Isabella Ippolito, Harmanpreet S. Sidhu, Emily Cacchione, Carlos Colangelo, Greg F. Slater, Rebecca T. Doyle

## Abstract

Rising fertilizer costs and dwindling freshwater supplies are driving interest in alternative agricultural amendments such as biosolids and reclaimed water. While these inputs can promote crop growth, they also contain persistent contaminants like PFAS, which may affect plants directly or indirectly via changes to plant-associated microbiomes. The extent and mechanisms of these effects remain poorly understood, particularly across crops with differing functional traits. We conducted two greenhouse experiments to evaluate the direct and microbially mediated effects of biosolid and reclaimed water amendments spiked with increasing contaminant concentrations on lettuce (*Lactuca sativa*), radish (*Raphanus sativus*), and green pea (*Pisum sativum*). Peas exhibited pronounced negative responses to biosolids, including reduced germination, survival, and biomass, which scaled with contaminant concentration and coincided with visible pathogen infection. Lettuce and radish showed minimal impact, suggesting that plant functional traits—such as symbiotic capacity and nutrient-dependent immune regulation—may mediate susceptibility to contaminant-induced stress. Soil microbiome diversity and composition were altered in a crop- and amendment-specific manner, but microbiomes shaped by prior exposure had limited effects on pea performance when transferred to new plants in the absence of ongoing stressors. These findings suggest that current soil conditions and active plant–microbiome interactions may outweigh legacy microbiome effects. Overall, our study provides a novel framework for evaluating the risks and benefits of wastewater-derived amendments, emphasizing the importance of functional trait variation in predicting plant responses to contaminants and guiding sustainable agricultural practices.

## Introduction

The soil surrounding a plant’s roots, known as the rhizosphere, is home to one of the most diverse communities on Earth – the soil microbiome (Fierer and Jackson 2006). Microbes within the soil microbiome play important roles in nutrient cycling, pathogen suppression, and promoting plant growth, stress tolerance, and disease resistance (Dubey et al. 2019; Philippot et al. 2024). Thus, soil microbiomes represent critical determinants of plant health (Banerjee and van der Heijden 2023), yet their composition and function are influenced by the surrounding environment, including the plant species present (Haichar et al. 2014). Agricultural amendments, while intended to boost crop yields, can unintentionally disrupt these microbiomes. For example, the addition of nitrogen-based fertilizers can shift microbial dynamics, increasing the abundance of less beneficial nitrogen-fixing bacteria (i.e., rhizobia) in legume-associated soils (Weese et al. 2015; Klinger et al. 2016). Given the central role of soil microbes in regulating plant health, understanding how microbiomes respond to environmental inputs, particularly agricultural amendments, and whether microbial responses impact crop performance, will be critical for optimizing amendment strategies that support both crop productivity and soil health.

Biosolids and reclaimed water, byproducts of wastewater treatment that offer nutrient-rich, water-conserving solutions for agriculture, are among the most widely promoted alternatives to conventional fertilizers and irrigation (Chen et al. 2013; Sharma et al. 2017). Although they offer agronomic benefits, these amendments often contain persistent contaminants, including polyfluoroalkyl substances (PFASs), perfluorooctane sulfonic acid (PFOSs), polybrominated diphenyl ethers (PBDEs), and antibiotics (Wu et al. 2015; Maddela et al. 2022). While the direct exposure effects of these contaminants on plant traits, such as toxicity and bioaccumulation, are increasingly recognized (Lillenberg et al. 2010; K. Kipper et al. 2010; Eggen et al. 2011; Sidhu et al. 2019; Krupka et al. 2024), their indirect, microbially-mediated impacts remain poorly understood. Contaminants may shift microbiome composition in ways that either buffer or intensify their effects on plants (Lau and terHorst 2020; Petipas et al. 2021; Angulo et al. 2022). For example, PFAS were found to reduce soil microbiome diversity (Cao et al. 2022), which could compromise the microbiome’s plant-growth promoting functions – though this link remains largely untested. Despite the growing use of these amendments, relatively few studies have examined how they influence microbiome dynamics over time, or whether contaminants within these amendments affects the microbiome’s ability to support plant growth (Schlatter et al. 2019). Additionally, biosolids can contain their own microbes (Reid et al. 2025), which can alter the resident microbiome’s functional composition and directly affect plant performance – either through microbial interactions or by introducing new functional traits. Understanding these microbial responses is essential for evaluating whether the benefits of alternative amendments outweigh the risks posed by the contaminants and microbiomes they carry.

The extent to which microbiomes can buffer or exacerbate the effects of agricultural contaminants on crop performance depends on several interacting factors: how the microbiome itself responds to the contaminant, the concentration of the contaminant, and the degree of plant reliance on microbial partners. Contaminants such as antimicrobials may disrupt microbiome structure, often reducing diversity and potentially impairing plant growth-promoting functions (Müller et al. 2002; Bell et al. 2005). However, soil microbiomes are often resilient to disturbance (Jiao Shuo et al. 2019), with diversity and function recovering over time (Zielezny et al. 2006; Ding et al. 2014), suggesting that microbial mediated effects may be transient. In contrast, direct phytotoxic effects of contaminants may be more immediate and severe, potentially overwhelming any beneficial microbial contributions. For example, tetracycline exposure inhibited pea plant growth and inhibited wheat root elongation (Yang et al. 2010; Krupka et al. 2024), as well as caused pea leaf yellowing and death of root tissue in seedlings growing in tetracycline for 120h (Margas et al. 2019), despite evidence that intermediate concentrations of the antibiotic were associated with higher microbiome diversity, likely via the suppression of dominant taxa (Zheng et al. 2020). Additionally, crop species differ in their dependence on microbial partners; legumes, including peas, rely heavily on rhizobia as a source of nitrogen, while non-legumes, including radish and lettuce, are more reliant on external nutrient inputs (Mathesius 2022). Consequently, microbially mediated effects of contaminants may be more pronounced in leguminous crops (Angle 1998). Despite growing interest in these microbially-mediated dynamics, most studies have focused on a single crop species or contaminant level or used unrealistically high contaminant levels, limiting our understanding of how microbial responses and plant outcomes vary across species with different microbial dependencies in agricultural systems.

Here, we investigate how alternative amendments, biosolids and reclaimed water, spiked with increasing environmentally-relevant contaminant concentrations, affect the growth performance of three agriculturally relevant crop species: lettuce (*Lactuca sativa*), radish (*Raphanus sativus*), and green pea (*Pisum sativum*). Our study had three main objectives. First, we directly exposed each crop species to amendments and contaminants and quantified plant trait responses, including germination, survival, and biomass, testing whether increasing contaminant levels altered plant performance and whether these responses varied by crop species. We predicted that biosolids, being more nutrient-rich than reclaimed water, would enhance plant performance (Nicholson et al. 2018), but that their benefits would decline with increasing contaminants due to physiological stress or interference with growth processes (Wu et al. 2015). Second, we characterized soil microbiome diversity and composition one week post planting and harvest to assess temporal shifts related to amendment and contaminant exposure and test for associations between these shifts and plant performance. We predicted initial changes in microbiome structure driven by nutrient inputs and contaminant selection (Mossa et al. 2017), followed by increased resilience over time (Jiao Shuo et al. 2019), with temporal dynamics being dependent on crop species. Third, we assessed the legacy effects of amendment and contaminant exposure on the microbiome’s ability to support plant growth by inoculating pea plants with soil slurries containing microbiomes from the first experiment. We predicted that microbiomes from contaminated soils would exhibit reduced pea growth compared to those from unamended soils with no contaminants added. By disentangling the direct and microbially-mediated effects of amendments across multiple crop species and contaminant concentrations, our study provides critical insight into the ecological trade-offs of using biosolids and reclaimed water in agriculture—highlighting the importance of microbiome dynamics in shaping plant responses to environmental contaminants.

## Materials and Methods

### Focal crop species

We selected three crop species—lettuce (*Lactuca sativa* var. Buttercrunch), radish (*Raphanus sativus* var. Crunchy Royale), and green pea (*Pisum sativum*, variety unspecified)—to capture a range of growth forms and functional traits relevant to plant-microbe interactions. Lettuce and radish, both non-leguminous dicots, have relatively low microbial dependence, while peas, as legumes, rely on symbiotic nitrogen-fixing rhizobia. These differences allowed us to test whether microbial reliance influences sensitivity to amendments and contaminants. Additionally, the crops differ in biomass allocation: lettuce and peas primarily produce above-ground biomass, whereas radish performance is more closely tied to below-ground growth. These traits may influence exposure pathways and susceptibility to soil contaminants. Finally, all three species reach maturity within 33–45 days, enabling completion of the experiment within two months with minimal temporal lag. Lettuce and radish seeds were sourced from JNB Seed Inc. (Regina, Saskatchewan, Canada), while pea seeds of the same cultivar were obtained from a commercial supplier who requested anonymity.

### Direct exposure experiment

From July to August 2024, we conducted a greenhouse experiment at McMaster University (Hamilton, ON, Canada) using two amendment types: biosolids (BS) and reclaimed water (RW). Each amendment was spiked with three contaminant concentrations: spiking level (SL) 1, SL-2, and an un-spiked control SL-0 (**Supplementary Methods S1**). SL-1 reflected contaminant concentrations commonly reported in studies of biosolid and reclaimed water amendments (Stevens et al. 2003; Panthi et al. 2019; Wang et al. 2019; Ding et al. 2020; McGrath et al. 2020; Semerád et al. 2020; Thompson et al. 2022). SL-2 represented concentrations approximately 2 –10 times greater than SL-1, simulating high-end environmental loading rates (Stevens et al. 2003; US Environmental Protection Agency 2009) and capturing variability across amendments sources and batches. This design allows us to test whether elevated contaminant levels diminish the agronomic benefits of wastewater-derived amendments.

We exposed the resulting six treatments (RW-0, RW-1, RW-2, BS-0, BS-1, BS-2) to each of our three crop species (lettuce, radish, peas), each being replicated three times, for a total of 54 experimental units (i.e., pots: 2 amendments x 3 spiking levels x 3 crop species x 3 replicates). We additionally included three unamended control pots without additional contaminant spiking (control-0) for each crop species, for a total of 63 pots (3 controls x 3 crop species = 9 control pots + 54 treated pots). Finally, we included six plant-free/soil only controls using magenta boxes (autoclavable and stackable pots with a self-contained fertilizer compartment; Bioworld, Dublin, Ohio, USA). Three of these magenta boxes contained autoclaved garden soil only, whereas the other three magenta boxes contained garden soil amended with 1% unspiked biosolids (w/w) that was then autoclaved, allowing us to account for any microbes naturally present in the greenhouse.

Pots used for planting were made from 2-gallon buckets (Home Depot, Atlanta Georgia, United States). All pots were filled with ∼5.8 kg of garden soil as the potting substrate (Bella Terra, S. Boudrias, Quebec, Canada; see **Supplementary Methods S2** for nutrient breakdown). To allow for drainage, we modified each pot by drilling holes at the bottom centre and covering them with fiberglass mesh pads to prevent soil loss. To simulate soil fauna -- a critical biotic factor found in natural conditions -- 15 adult earthworms were added to each pot. To make up the biosolid amendments, soils were mixed with 1% (w/w) of biosolids, an application rate typical in agriculture (US Environmental Protection Agency 2009) using a concrete mixer to ensure homogeneity. To implement spiking treatments (SL1, SP2), contaminants were diluted to their appropriate concentrations (**Supplementary Methods S1**) using water mixed with 1% methanol or 1% acetone, added to a spray bottle, and then sprayed directly onto a thin layer of biosolids (Sidhu et al. 2019). Spiked biosolids were allowed to equilibrate for a week, allowing solvents to evaporate prior to mixing with the garden soil. For the reclaimed water treatments, we used reclaimed water that was left unspiked (SL-0) or spiked with the appropriate concentrations (**Supplementary Methods S1**) by adding contaminants directly to the reclaimed water prior to each irrigation event. Anaerobically digested and heat dried biosolids and tertiarily treated reclaimed water were provided by a local wastewater treatment plant in southern Ontario that serves over half a million people and treats an average of ∼410 million litres of wastewater daily.

To ensure comparable plant density and minimize variation in biomass accumulation due to crowding, we sowed 12 lettuce seeds, 20 radish seeds, and 25 pea seeds per pot. Plants were grown for 45-46 days (lettuce and peas) and 33 days (radish), reflecting typical growth cycles for each species under greenhouse conditions and allowing us to capture relevant developmental stages for trait assessment. To account for growth differences due to position in the greenhouse (i.e., temperature, light, humidity), pot positions were randomly assigned with respect to treatment and crop species across three benches and were re-randomized on a weekly basis until the end of the experiment. Throughout the experiment, all pots were watered to field capacity to ensure soil pores had an optimal balance of air and water. We watered soils every other day based on protocol detailed in Sidhu et al. (2019). Pots in the reclaimed water treatment were only watered with reclaimed water (plus their respective spiking levels), whereas for biosolid-amended pots and unamended controls, watering was completed using tap water from the greenhouse hose. Miracle Gro® (N:P:K of 24:8:16, plus micronutrients) was applied before seeding and every week after seedling emergence until plant harvesting. The total fertilizer dose added per bucket was equivalent to ∼195 kg N ha−1, ∼49 kg P ha-1, and 188 kg K ha-1.

We measured germination rates for all plants one week after sowing. Forty days post-planting, we assessed the Normalized Difference Vegetation Index (NDVI) for pea plants using a PSI Plant Pen (Qubit Systems, Kingston, Ontario, Canada). NDVI is a commonly used non-destructive proxy for leaf chlorophyll A content (Tucker 1979), providing an integrative indicator of leaf nitrogen content, often measured for legumes to assess symbiotic effectiveness. After calibrating the device, we randomly selected three leaves per pot for NDVI measurements. At harvest (day 46 for peas and 45 for lettuce, and 33 for radish), we gently removed plants from the soil and washed them sequentially in three tubs of clean tap water, replacing the water between pots to prevent cross-contamination.

During the experiment, we unexpectedly observed signs of pathogen infection in several pea plants amended with biosolids. We therefore counted the number of peas with visible infection to account for this. At harvest, we patted all plants dry and separated roots from shoots using sterilized scissors, recording both shoot and root fresh weight. Both shoots and roots were freeze-dried, and their dry weights were recorded. For peas only, we recorded the wet and dry pea pod weights, the number of pea pods and flowers per pot, and noted whether any nodules were present -- the root structures containing rhizobia. In total only three of 21 pots had pea plants with nodules (6 – 13 nodules per plant). All nodules were green, meaning they were likely inactive for nitrogen-fixation.

### Soil microbiome sequencing

To characterize the soil microbiome over the course of the experiment, we collected soil samples at two timepoints: one week after planting and at harvest (33 - 46 days post-planting, depending on crop species), for a total of 126 experimental samples (63 pots x 2 timepoints) plus the six soil-only controls. From each pot, we sampled ∼10 g of rhizospheric soil from five random points near the roots, combined the subsamples, thoroughly homogenized them to reduce within-pot heterogeneity, and stored the composite samples at –20 °C. Once all samples were collected from the two timepoints, we added ∼200 mg of each to 2 mL bead tubes pre-filled with an extraction buffer made of 800μl of 200mM monobasic NaPO_4_ (pH 8) and 100μl of Germanium Diselenide (GES). Samples were submitted for DNA extraction and 16S rRNA amplicon sequencing targeting the V4 region, which provides high taxonomic resolution for soil-associated microbial taxa, down to the genus level (Wasimuddin et al. 2020; Acharya et al. 2021). Sequencing was performed at the Surette Lab, McMaster University ([link](https://genomics.healthsci.mcmaster.ca/services/next-generation-sequencing/amplicon-sequencing/)). In addition to experimental samples, we submitted 24 “source” samples that included triplicates of each of the following: biosolids, reclaimed water – each at the three spiking levels -- garden soil (unautoclaved), and tap water, allowing us to characterize microbiomes present in these sources. For aqueous samples (tap and reclaimed water), we added 600 µL to each extraction tube and processed them using the same protocol as the solid samples.

### Read processing with DADA2

Cutadapt was used to filter and trim adapter sequences and PCR primers from the raw reads with a minimum quality score of 30 and a minimum read length of 100bp (DOI:10.14806/ej.17.1.200). Sequence variants were then resolved from the trimmed raw reads using DADA2, an accurate sample inference pipeline for 16S amplicon data (doi: 10.1038/nmeth.3869). DNA sequence reads were filtered and trimmed based on the quality of the reads for each Illumina run separately, error rates estimated, and sequence variants determined via DADA2. Sequence variant tables were merged to combine all information from separate Illumina runs. Chimera sequences were removed, and taxonomy was assigned using the SILVA database version 1.2.8 (Quast et al. 2013; Yilmaz et al. 2014; Glöckner et al. 2017).

### Microbiome legacy effects experiment

From October to November 2024, we conducted a follow-up greenhouse experiment to assess the legacy effects of soil microbiomes previously exposed to amendments and varying contaminant concentrations on pea plant growth. We chose pea as our focal species for this experiment because we *a priori* expected it to be more reliant on microbes for growth, and from the direct exposure experiment, showed the most significant responses to amendments and contaminants compared to the other two species. We sowed 73 green pea seeds (same variety used in our first experiment) into sterilized Cone-tainers™ (RL98K10R; Tangent, Oregon, United States) filled with a 1:1 mixture of autoclaved peat moss/perlite mix (Berger mix no. 6, HP; Saint-Modeste, Quebec, Canada) and Turface (MVP; Buffalo Grove, Illinois, United States). Using this low-nutrient, sterile potting mixture rather than the garden soil used in the previous experiment allowed us to assess the ability of microbiome to support plant growth without the additional effects of nutrients and other microbes in the soil. Each Cone-tainer was fitted with an autoclaved cotton wick and placed above a sterile 50 mL centrifuge tube to collect flow-through. Prior to planting, we surface-sterilized pea seeds by submerging them in 0.7% hypochlorite for 30 minutes using a metal tea ball, followed by five rinses with deionized water. One seed was sown per Cone-tainer.

To maintain consistent nutrient conditions throughout the experiment, we saturated the soil with 0.625 mM nitrogen fertilizer and watered daily until germination. Once seedlings had germinated, we inoculated them with soil slurries prepared from samples collected at the end of the previous greenhouse experiment. Soil slurries were made by mixing 8 grams of soil with 40 mLs autoclaved physiological saline solution (0.85% w/v NaCl2 in water; Weese et al. 2015). Slurries were shaken by hand for one minute, allowed to settle, and the supernatant was transferred to a new tube and settled again. We pipetted 10 mL of the resulting slurry onto each of three replicate pea plants per treatment. There were 21 soil slurries in total: 18 from the experimental treatment pots (2 amendments x 3 contaminant levels x 3 replicates) and three from the unamended, uncontaminated control pots, resulting in 63 plant replicates in total (21 pots x 3 replicates). In addition to the soil-slurry inoculated plants, the seven remaining germinated seedlings were inoculated with sterile saline to serve as controls, for a total of 70 plants.

We inoculated pea plants 21 days post-planting, after confirming sufficient germination, to avoid wasting limited soil slurry volumes on seeds that failed to germinate. While this approach prevented assessment of early-stage traits such as germination, our primary focus was on later-stage performance metrics (e.g., biomass). Post-emergence inoculation ensured that all plants began the experiment from a consistent developmental stage, reducing variability due to uneven germination and allowing us to isolate the effects of microbiome treatments on growth. Throughout the experiment, we measured plant height and NDVI values weekly. Height was measured from the soil surface to the shoot tip using a measuring tape. NDVI values were collected using a PSI Plant Pen (Qubit Systems, Kingston, Ontario, Canada), calibrated prior to use, with three randomly selected leaves measured per plant. Prior to harvest, we recorded leaf counts and flower counts for any flowering plants. Nodule counts were recorded during dissection. After 45 days of growth, consistent with the first experiment, we performed a full destructive harvest. We recorded shoot fresh weight, then dried shoots in paper coin envelopes at 50 °C for three days. Roots were rinsed in tap water, dried on paper towels, and stored in 50 mL centrifuge tubes with silica gel and cotton until nodules were dissected. Nodules present on roots were removed and counted. We recorded root fresh weight and then dried roots at 50 °C for three days before measuring dry weights.

### Statistical analyses: crop performance

We used a linear modeling framework in R (v4.5.1; R Core Team, 2025) to assess the direct and indirect effects of amendment type and contaminant spiking level on plant traits, implementing a two-part modeling strategy. First, we tested whether amendment type (biosolids, reclaimed water, unamended control) influenced plant traits in the absence of spiked contaminants. This model included the unamended control as a reference and allowed us to test for crop species × amendment type interactions, capturing species-specific responses to the amendments themselves. Second, we evaluated the effects of contaminant spiking level (coded as an ordered factor: 0, 1, 2) within each amendment type. Because contaminant concentrations were not directly comparable between biosolids and reclaimed water, we ran separate models for each amendment type, excluding the unamended controls. These models tested for dosage-dependent effects of contaminants on crop performance as well as whether these effects varied by crop species.

For the direct exposure experiment (Experiment 1), response variables included germination rate (total seeds germinated/total seeds planted), survival rate (total plants harvested/total seeds planted, dry total biomass (dry root + shoot biomass), per plant biomass (dry total biomass/ total plants harvested), shoot moisture content ([wet - dry shoot biomass]/ dry shoot biomass), and shoot-to-root ratio (dry shoot/dry root biomass), all measured at the pot-level. For peas, we additionally analyzed dry pod biomass, per pod biomass (dry pod biomass / total pods), flower count, pod count, NDVI, and infection rate (number of peas visibly infected by pathogen / total seeds planted), all measured at the pot-level. In addition to linear models, we conducted a linear regression using the base::lm() R function to test whether pea survival was correlated with infection rate.

For the microbiome legacy effects experiment, we first used the emmeans package (v1.11.1; Lenth, 2025) to calculate the estimated marginal means (EM means) of response variables measured for each of the 21 soil slurries constructed from replicate pots in Experiment 1, plus saline-inoculated controls (N = 3). Response variables included plant height, NDVI, and leaf number, each measured weekly throughout the experiment, as well as dry total biomass, shoot moisture, and shoot-to-root ratio measured at harvest, all measured at the plant-level, as only one seed was sown per pot. We then treated these EM means as response variables in models testing the same two hypotheses described for Experiment 1.

To test the significance of interaction terms in all models, we used type III sums of squares from the *car* package (v3.1.; Fox & Weisberg, 2019). Effect sizes and 95% confidence intervals were calculated using Dunnett-style contrasts (for models with unamended controls as reference) and polynomial contrasts (for contaminant level models), both implemented in the *emmeans* package. The *emmeans* package automatically applies the most appropriate p-value adjustment for multiple comparisons, and these adjusted values are reported. To meet ANOVA assumptions, we log-transformed all non-count data, while count-based traits (e.g., pod, flower, leaf number) were modeled using a Poisson distribution with a log link function. Model assumptions were verified by visually inspecting residual diagnostics plots.

### 16S microbiome analyses

We used the phyloseq package in R (v1.50.0; McMurdie and Holmes 2013) to create a phyloseq object integrating amplicon sequence variant (ASV) counts, taxonomic classifications (Kingdom to Species), and sample metadata (pot ID, amendment/contaminant treatment, crop species, and timepoint). We removed ASVs assigned to non-bacterial kingdoms and any host mitochondrial DNA prior to analysis.

#### Alpha Diversity

To control for differences in sequencing depth, all samples were rarified to 11,934 reads—the minimum read count after quality filtering—using rarefy_even_depth() function in *phyloseq* (v1.52.0; McMurdie and Holmes 2013). Shannon and Inverse Simpson (1/*D*) diversity indices were calculated for each sample using estimate_richness(). The Shannon index reflects richness and evenness, while Inverse Simpson index emphasizes dominance and evenness (Roswell et al. 2021). These metrics served as response variables in linear mixed models (LMMs) similar to LMs described above, with two key differences: (1) Models included crop species, amendment type (biosolids, reclaimed water, unamended control) or contamination level (SL-0, SL-1, SL-2), sampling timepoint (one week post planting, harvest), and their respective two- and three-way interactions. (2) a random effect of pot was added to account for repeated measures. This approach allowed us to assess amendment and contaminant effects, as well as changes in microbiome diversity over plant development and whether crop identity modulated these effects. We used type III sums of squares to assess the significance of all interaction terms and similarly used the emmeans package to estimate contrasts similar to the first set of models. To verify ANOVA assumptions were met, Shannon and Inverse Simpson indices were log-transformed prior to modeling, and residual diagnostics were visually inspected.

#### Beta Diversity

We assessed microbial community composition using Bray-Curtis dissimilarity from unrarefied ASV abundances (results were similar with rarefied data). Community structure was visualized using Principal Coordinates Analysis (PCoA) and Non-Metric Multidimensional Scaling (NMDS) via the *phyloseq*::ordinate() function, including both source and experimental samples. To test the effects of crop species (three levels), treatment (combined amendment type and spiking level, seven levels), and timepoint (two levels) on composition, we used Constrained Analysis of Principal Coordinates (CAP) via the *phyloseq*::capscale() function with Bray-Curtis distances. Separate CAP models were additionally run for each timepoint (1-week post-planting and harvest) to assess temporal shifts in explanatory power for both crop species and soil treatment. To test whether plant performance traits were associated with microbiome composition, we initially used a forward-selection approach to identify continuous predictors that significantly improved explanatory power, using these predictors in CAP models. Traits included dry total biomass, shoot:root ratio, shoot moisture, and survival for all crop species, with additional pea-specific traits (dry pod biomass, and pod/flower number). Trait–microbiome associations were assessed using the envfit() function with 999 permutations.

## Results

### Crop-dependent responses to amendments and contaminants

We first tested whether soil amendments (biosolids [BS] and reclaimed water [RW], without added contaminants) influence plant performance relative to unamended controls. Amendments had minimal effects on biomass traits (**Fig. 1; Supp. Table S1 & Fig. S1A**), but significantly reduced germination (F [2, 18] = 4.61, p = 0.0241) and survival (F [2, 18] = 4.51, p = 0.0259), regardless of crop species (**Supp. Table S1**). These effects were most pronounced in peas, where survival was significantly reduced under BS amendment compared to unamended controls (**Supp. Table S1 & Fig. S1A**).

**Figure 1:**
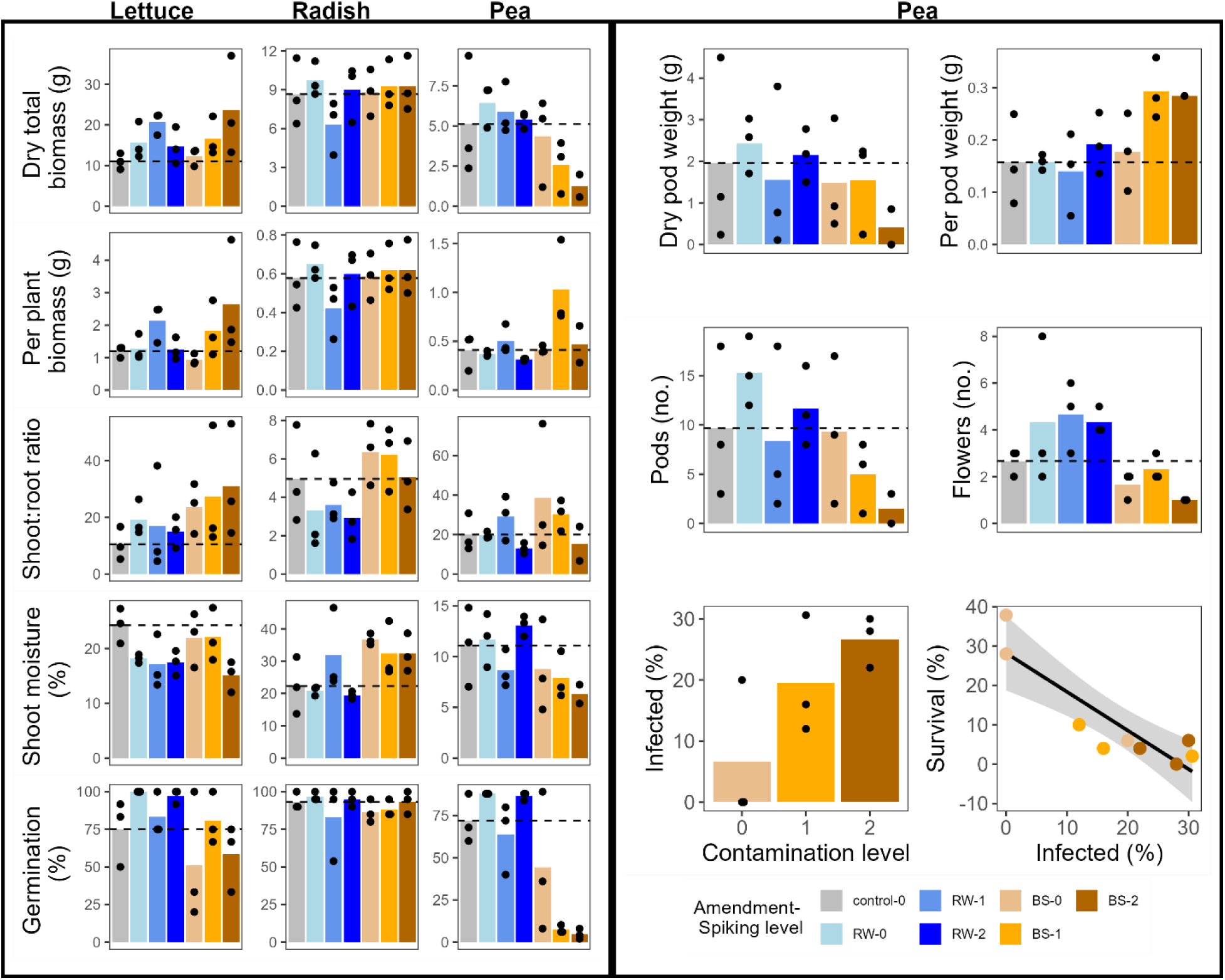
Crop species differ in their responses to amendments and contaminant concentrations. **Left,** Trait responses across crop species under different amendment and contaminant treatments. Bars show raw means; points represent individual pot replicates (N = 3 per treatment). Dashed lines indicate the mean of unamended controls without contaminants. **Right,** Trait responses in peas only. Bottom row shows correlations between traits and contaminant levels in biosolid-amended peas (N = 3 per spiking level). *Key take-away: Peas exhibited stronger stress responses to biosolid amendments and contaminants than lettuce or radish, highlighting crop-specific sensitivity linked to functional traits*.

Next, we tested whether increasing contaminant concentrations affected plant performance within each amendment type. Responses varied by amendment and crop species (**Fig. 1**; **Supp. Table S2**, **Supp. Fig. S1B**). Under RW, most traits showed non-linear responses to contaminant level (i.e., significant quadratic contrasts). Germination (F [2, 18] = 5.23, p = 0.0162) and survival (F [2, 18] = 3.89, p = 0.0393) declined most at intermediate concentrations (i.e., positive quadratic terms). For biomass (per plant) and shoot moisture, effects depended on crop species (biomass: F [4, 18] = 4.2, p = 0.0142; shoot moisture: F [4, 18] = 4.22, p = 0.0139). Lettuce and peas had higher biomass at intermediate levels (i.e., negative quadratic terms), whereas radish had lower biomass. Radish also had higher shoot moisture at intermediate levels, while peas showed the opposite pattern (**Supp. Table S2**, **Supp. Fig. S1B**). Under BS, responses to contaminant level were more linear: biomass (per plant) increased with contaminant level (F [2, 17] = 4.69, p = 0.0239), especially in lettuce. However, contaminant effects on germination (F [4, 18] = 4.3, p = 0.0129) and survival (F [4, 18] = 3.08, p = 0.0425) varied by crop species, with peas mostly negatively affected as levels increased. Pea survival was strongly linked to pathogen infection (t = −5.1, p = 0.00139; Slope ± SE = −0.983 ± 0.192), which increased with contaminant level (t = 2.67, df = 6, p = 0.0368; linear est. [95% CI] = 2.3 [0.195 to 4.4]; **Fig. 1**). Overall, RW had mixed effects, while BS mainly harmed peas at higher contaminant levels, likely due to pathogen susceptibility.

### Soil microbiome responses to amendment and contaminant exposure

We hypothesized that amendment type would alter soil microbiome diversity, with higher contaminant concentrations leading to reduced diversity. Diversity was strongly driven by sampling timepoint (Shannon: χ²[1] = 12.2; Inverse Simpson: χ²[1] = 23.2; both p < 0.001), with values significantly lower one-week post-planting than at harvest (**Fig. 2A**; **Supp. Table S3**; **Supp. Fig. S2A**). For Shannon diversity, this temporal effect varied by crop species (species × timepoint: χ²[2] = 6.84, p = 0.033) and amendment type (species × amendment × timepoint: χ²[4] = 14.2, p = 0.0066), whereas Inverse Simpson showed no significant interactions (**Supp. Fig. S2B**). At one-week post planting, unamended pea diversity was significantly lower than lettuce and radish, but these species differences disappeared by harvest (**Supp. Table S3; Supp. Fig. S2B**). RW amendment early on increased pea microbiome diversity while reducing lettuce diversity (**Supp. Table S3**); however, differences among crop species diminished by harvest (**Supp. Fig. S3A**). Overall, microbiome diversity declined initially but by harvest, recovered across all species, highlighting resilience in community structure despite early amendment-driven shifts. Testing contaminant concentration effects within BS and RW amendments revealed that timepoint remained the dominant factor (p < 0.01), alongside species and species × timepoint interactions (**Fig. 2A**: **Supp. Table S4**; **Supp. Fig. S2A**). Under RW amendment one-week post planting, lettuce diversity increased modestly with contaminants, but effects were negligible by harvest (**Supp. Table S4; Supp. Fig. S3B**). Overall, contaminant concentration had minimal influence compared to species and timepoint, indicating that temporal and host factors were the primary determinants of microbiome diversity.

**Figure 2:**
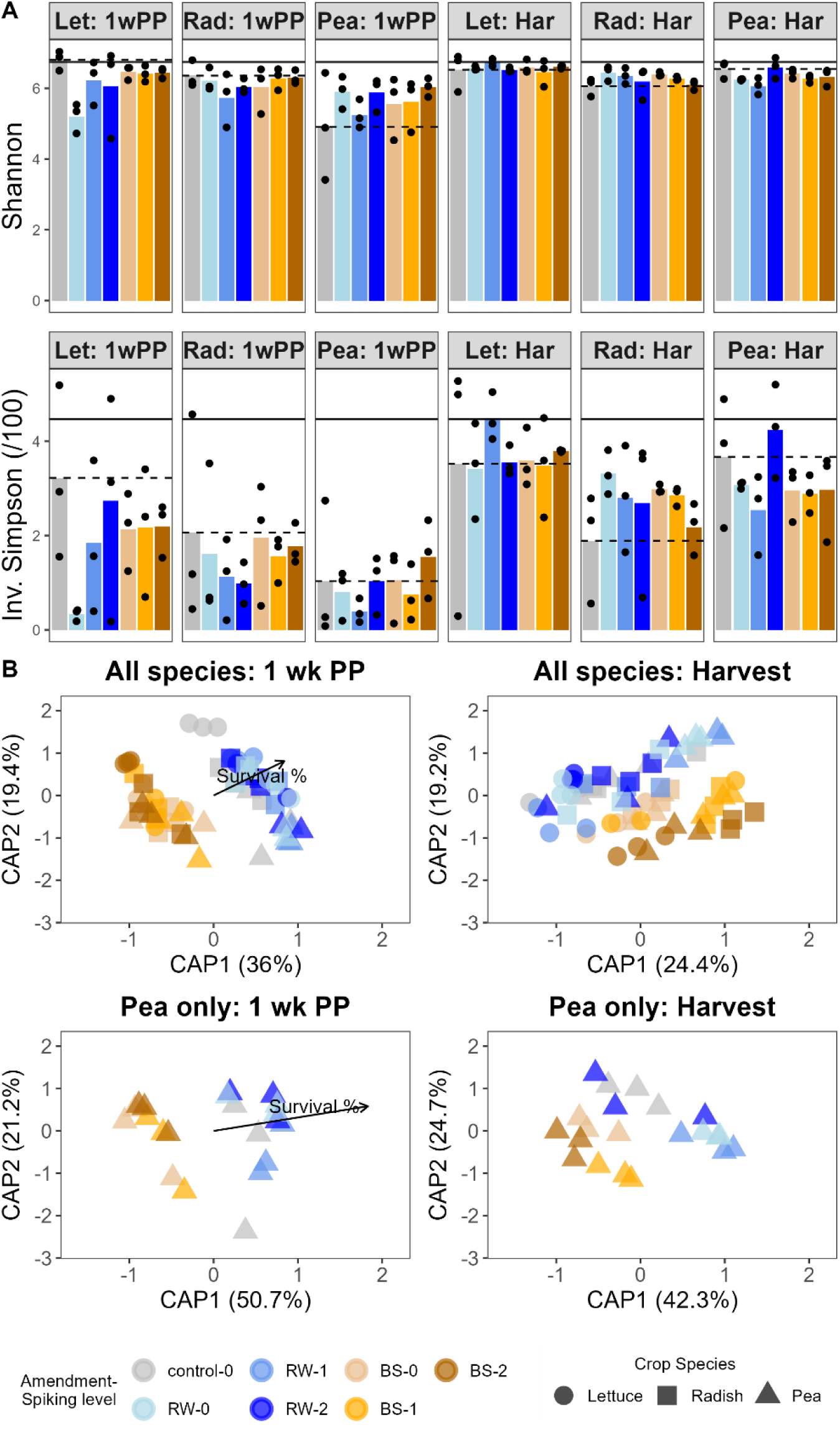
Microbiome diversity and composition are shaped by both crop species and soil treatment. **A** Alpha diversity (Shannon and Inverse Simpson) of soil microbiomes sampled one-week post-planting (1wkPP) and at harvest (Har) across crops (N = 3 per treatment). Dashed lines = unamended controls; solid lines = initial garden soil. **B** Constrained Analysis of Principle Coordinates (CAP), a form of Distance-based Redundancy analysis (dbRDA), with ordination of microbiomes across crop species and just for peas showing variation explained by soil treatments. Arrows indicate traits significantly associated with community composition (p < 0.05). *Key take-away: Soil treatments altered microbiome diversity and composition, especially in peas, but diversity tended to recover over time—suggesting microbial resilience that may buffer long-term plant impacts*.

Finally, we assessed the relative contributions of crop species and soil treatment (amendment + contaminant level) on microbiome composition using Bray-Curtis dissimilarity. An initial examination of the PCoA ordination revealed clear clustering by timepoint and treatment, indicating strong temporal and amendment-driven effects on community structure (**Supp. Fig. S4**). Both amendment and species terms were significant in CAP models (i.e., species x treatment interaction: F [12, 79] =1.46, p = 0.001; **Fig. 2B**; **Supp. Table S5**), but their relative importance shifted over time: crop species explained 10% of the variation initially and 13% at harvest, while soil treatment declined from 27% to 21%. Associations between microbiome composition and plant traits weakened over time (**Supp. Table S5**). Early in the experiment, microbiomes across all three species were linked to survival (**Fig. 2B**), primarily driven by pea (**Supp. Fig S5**). By harvest, these associations were largely absent (**Supp. Table S5**). This pattern suggests that while soil treatment and crop species shape microbiome composition, their influence is dynamic, and early microbial interactions may mediate crop-specific responses to stressors.

### Microbiome legacy effects were minimal on pea performance

To isolate the legacy effects of microbiomes shaped by soil amendments and contaminants, we inoculated a new cohort of peas with microbiomes harvested from the direct exposure experiment, without reapplying any amendments or contaminants. This approach allowed us to assess how amendment- and contaminant-induced shifts in the microbiome alone influenced pea performance, independent of direct chemical exposure. Overall, microbiome legacy effects had limited influence on pea traits, except for shoot-to-root ratio, which was significantly impacted by prior amendment type (F[2, 6] = 13.8, p = 0.00565; **Fig. 3**). Microbiomes from BS-amended soils tended to increase shoot-to-root ratio (BS vs. control: t = 2.82, df = 6, p = 0.0553; log₂FC [95% CI] = 0.519 [–0.0144 to 1.05]), while those from RW-treated soils showed a trend toward reducing this trait (RW vs. control: t = –2.44, df = 6, p = 0.0917; log₂FC [95% CI] = –0.448 [–0.981 to 0.0857]). For microbiomes previously exposed to increasing contaminant concentrations within BS and RW amendments, we found a linear decline in shoot-to-root ratio (t = –4.8, df = 6, p = 0.003; slope = –0.551 [–0.831 to –0.27]) only under BS-amendment (**Fig. 3C**). These results indicate that while direct exposure to amendments and contaminants significantly impact microbiome diversity and composition, such indirect effects on plant traits are relatively modest when microbiomes are transferred to a new host, likely due to microbiome resiliency. In the context of our broader findings, this suggests that current microbiome diversity and composition—shaped by both crop species and soil treatment—may play a more dominant role in influencing plant performance than legacy effects alone.

**Figure 3:**
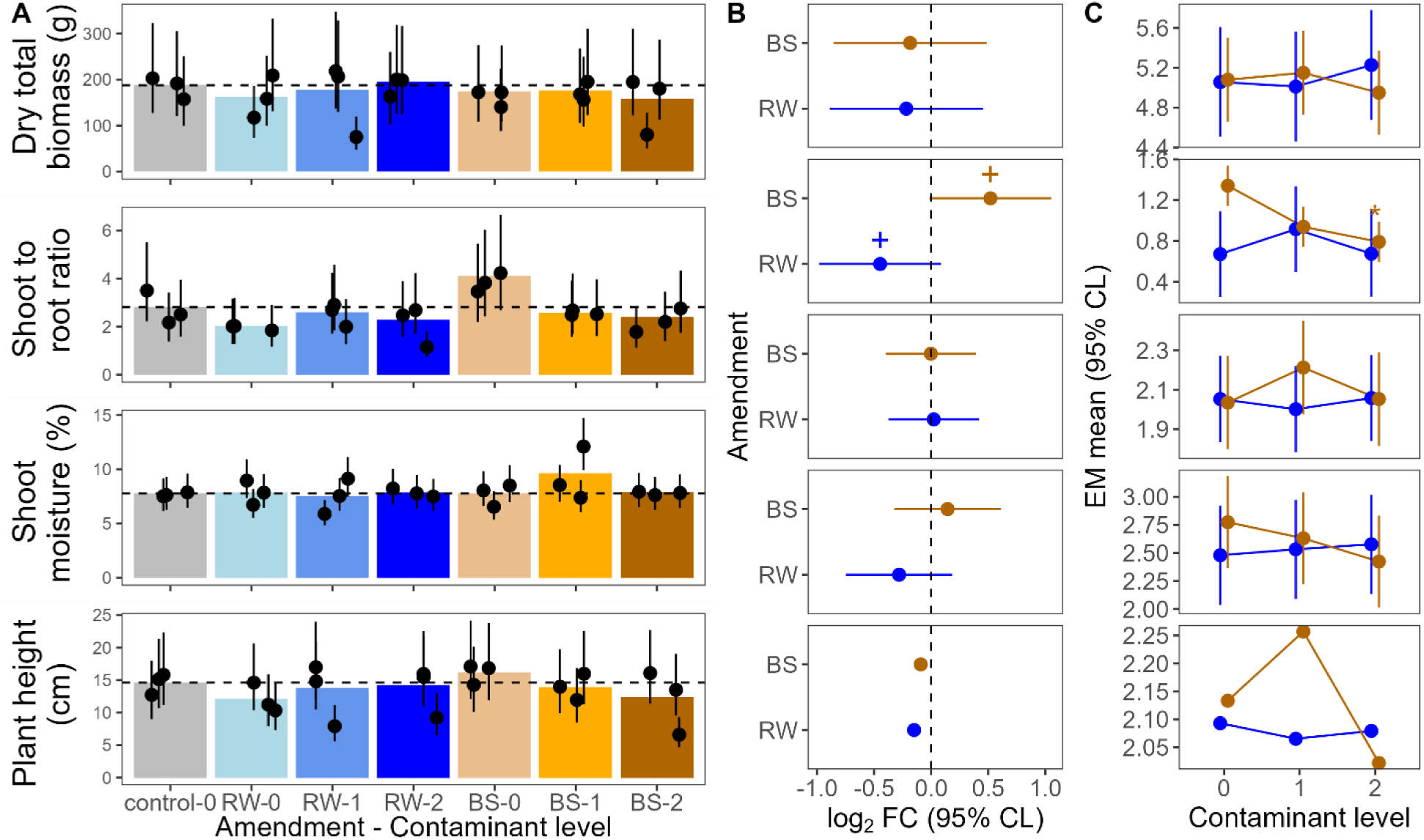
Microbiome legacy effects on pea performance are weaker in magnitude compared to direct exposure effects. **A** Trait response in peas inoculated with legacy microbiomes. Bars show raw means; points represent estimated marginal means (N = 3 per slurry, 9 per treatment). Dashed lines = unamended controls from direct exposure experiment. **B** Dunnett-style contrasts showing log2 fold changes in traits for biosolid (BS) and reclaimed water (RW) treatments vs. controls (spiking level 0). **C** Linear and quadratic trait responses to increasing contamination levels (spiking 0–2) for each amendment type. Points = estimated marginal means; error bars = 95% confidence limits. Asterisks (*) and plus signs (+) indicate significance (p < 0.05 and 0.05 < p < 0.1, respectively). *Key take-away: Legacy microbiomes had limited effects on pea performance in the absence of contaminants, suggesting that current soil conditions and microbial recovery may outweigh historical microbial legacies*.

## Discussion

As agriculture increasingly turns to alternative soil amendments, including biosolids and reclaimed water, understanding how these inputs influence crop performance—both directly and through their effects on soil microbiomes—is essential for predicting their suitability across diverse cropping systems. To address this gap, we assessed the direct effects of amendments and contaminants on multiple crop species, as well as the legacy effects of microbiomes shaped by those treatments on a new cohort of peas. Our findings reveal three key insights. First, crop responses to biosolid and reclaimed water amendments are not only species-specific but also functionally driven; peas, as legumes, showed heightened pathogen susceptibility that scaled with contaminant concentration, a pattern not observed in non-legumes. Second, amendment-induced shifts in microbiome diversity and composition can persist over time, but their relevance to plant performance is crop-dependent. Third, microbiomes shaped by past exposures have limited influence on plant growth in the absence of ongoing stressors, suggesting that current soil conditions and plant–microbiome interactions may play a more dominant role than microbial legacies alone. Together, these insights underscore the importance of considering crop-specific functional traits when evaluating the ecological and agronomic consequences of alternative soil amendments.

### Pea-specific pathogen susceptibility under biosolid and contaminant stress

Both biosolids and reclaimed water can have variable effects on plant traits, from beneficial to harmful (Moreno et al. 1997; Lillenberg et al. 2010; K. Kipper et al. 2010; Eggen et al. 2011; Sridhar et al. 2014), yet most studies focus on single crop. By growing lettuce, radish, and peas in the same experiment, we compared trait responses across species. While lettuce and radish showed mostly neutral or positive responses, peas exhibited pronounced negative effects when grown in biosolid-amended soil, particularly during early developmental stages such as germination. These early effects cascaded to affect growth, reproduction, and survival.

Visible infection occurred only in peas grown with biosolids, with severity increasing at higher contaminant levels. This species- and amendment-specific pattern underscores a key interaction between crop identity, amendment type, and contaminant load, though mechanisms remain unclear. Contaminants may have compromised pea immune defenses, increasing vulnerability to a pathogen present in biosolids, or the association may reflect correlated environmental conditions rather than direct contaminant effects. Infection was absent in reclaimed water treatments and in lettuce and radish treatments, pointing to a pea-specific pathogen introduced or favoured by biosolid properties.

Previous studies report contaminant-driven growth effects in peas (e.g., titanium dioxide nanoparticles, perfluoroalkyl acids; Giorgetti et al. 2019, Blaine et al. 2014; Alaoui et al. 2025), but none assessed disease incidence as a function of contamination levels. Although biosolids generally contain higher overall contaminant loads compared to reclaimed water (Dennis et al. 2025), many compounds are strongly sorbed and less bioavailable than the more mobile contaminants in reclaimed water (Borgman and Chefetz 2013), supporting the hypothesis that biosolids acted as the pathogen source rather than contaminants being the primary driver of infection.

Peas, as legumes, may also be predisposed to infection due to their reliance on symbiosis with nitrogen-fixing rhizobia. Under high nitrogen conditions, such as the nutrient-rich soil and weekly NPK fertilization used in our experiment, legumes typically reduce investment in symbiosis in favour of direct nitrogen uptake (Reid et al. 2011). Indeed, we observed few nodules on pea roots at the end of the direct exposure experiment. Reduced symbiosis and elevated nitrogen availability have been linked to increased pathogen susceptibility in legumes (Rey et al. 2013; Gourion et al. 2015; Thalineau et al. 2018; Zitnick-Anderson et al. 2021; Laine 2023). For example, nitrogen supplementation in *Medicago truncatula* heightened susceptibility to *Aphanomyces euteiches*, a major pathogen of field-grown peas (Thalineau et al. 2018; Zitnick-Anderson et al. 2021), and disruption of Nod factor receptor genes, which are required for symbiosis establishment with rhizobia, heightened vulnerability to both *A. euteiches* and *Colletotrichum trifolii* (Rey et al. 2013; Gourion et al. 2015). Other soil factors, such as pH, iron, and potassium, can influence pea disease outcomes (Oyarzun et al. 1998; Zitnick-Anderson et al. 2020), but the absence of infection in lettuce and radish suggests host-specific susceptibility rather than exposure alone.

Although direct evidence linking biosolids to pea pathogen susceptibility is lacking, biosolids can alter microbial communities and soil suppressiveness in ways that affect plant–pathogen interactions (Sharma and Reynnells 2016; Marie et al. 2024). Shifts toward stress-tolerant microbial taxa or changes in nutrient availability could compromise plant immune responses or favour pathogen proliferation (Mur et al. 2017; Sharma and Reynnells 2016; Reid et al. 2025). These mechanisms, observed in other crops and amendment types, warrant further investigation in legumes, which may be particularly sensitive to microbial and chemical disruptions due to their reliance on symbiotic nitrogen fixation.

Taken together, our findings suggest that biosolid amendment contributed to pea pathogen susceptibility through interacting factors -- contaminant-induced stress, nitrogen-mediated immune regulation, and crop-specific pathogen proliferation in biosolid-enriched environments. While the exact mechanism and pathogen identity remain unknown, these results underscore the importance of considering plant functional traits—such as symbiotic capacity and nutrient acquisition strategies—when evaluating the risks and benefits of alternative wastewater-derived amendments. Future work should disentangle these interactions to better predict amendment outcomes across crop types.

### Crop-specific associations between microbiome shifts and plant traits

Soil microbial communities play a central role in nutrient cycling, plant health, and stress resilience (Dubey et al. 2019; Banerjee and van der Heijden 2023; Philippot et al. 2024). Understanding whether amendment- and contaminant-induced shifts in crop-associated microbiomes influence plant trait is critical, especially because these associations may vary across crop species with distinct functional traits. By sampling soils from different crops grown under identical amendment and contaminant conditions at two timepoints, our study allowed us to disentangle how crop identity interacts with soil treatments to shape microbiome–plant trait relationships, and whether these associations shift over time.

Temporal sampling revealed that although the influence of soil treatments on microbiome composition diminished over time, it remained substantial even at harvest, exceeding the explanatory power of crop species. Notably, microbiome diversity—especially when measured using Inverse Simpson—was initially reduced but tended to recover by harvest, with pea-associated microbiomes “catching up” the most. This recovery suggests a degree of microbiome resilience, which may buffer long-term impacts on plant performance and help explain limited legacy effects in the follow-up experiment (discussed more below).

Despite persistent microbial shifts, their functional relevance appeared crop-specific. In peas, early microbiome composition was associated with survival, a pattern largely absent in lettuce and radish. These pea-specific associations suggest greater sensitivity to microbial perturbations or a stronger reliance on microbial partners in this legume species. Early disruptions may trigger stress responses that persist through development, ultimately reducing yield (Liu et al. 2025). However, the direction of causality remains unclear. Microbiome shifts may drive plant stress, or stressed plants may alter microbiomes (Müller et al. 2002; Bell et al. 2005; Angulo et al. 2022). For example, in biosolid-amended soils with high contaminant concentrations, where few peas survived, reduced root exudates could have lowered microbiome diversity rather than microbial shifts causing poor survival. While our data cannot confirm directionality, these findings highlight the potential importance of early microbiome dynamics in shaping long-term plant outcomes—an aspect often overlooked in studies that sample at a single timepoint.

Overall, our results demonstrate that microbiome shifts did not consistently translate into trait changes (Han et al. 2025). Instead, their effects were context-dependent, shaped predominantly by crop identity and timing. Importantly, amendment impacts extend beyond chemical composition to include lasting alterations in soil microbiomes, which may interact with crop physiology in species-specific ways. Future experiments that manipulate microbiomes independently of amendments across plant developmental stages will be key to disentangling these interactions and clarifying whether early microbiome disruptions exert more lasting effects than later ones.

### Limited legacy effects of microbiomes shaped by past soil treatments

Understanding how past soil conditions influence current plant performance is critical in agricultural systems, where crop rotations and historical management practices shape the soil microbiome long after amendments or contaminants are no longer present (White 1970; Dias et al. 2015; Yang et al. 2023). While many studies have focused on the immediate impacts of soil amendments on plant–microbiome interactions (Wang Zhonghua et al. 2022; Yu Yitian et al. 2024), few have tested whether microbiomes shaped by past exposures continue to influence plant traits in the absence of ongoing chemical inputs. Our approach—transferring microbiomes from soils previously exposed to amendments and contaminants onto a new cohort of peas grown in untreated soil—allowed us to isolate these legacy effects.

Despite clear and persistent shifts in microbiome diversity and composition in the original (direct exposure) experiment, legacy microbiomes had relatively modest effects on pea performance when contaminants and amendments were no longer present. These modest legacy effects may reflect the resilience of microbial communities observed earlier, where microbiome diversity—initially reduced, especially in peas—rebounded over time. Such recovery could restore key microbial functions and reduce the likelihood of persistent impacts on plant traits in subsequent generations (Jiao Shuo et al. 2019). These results suggest that current soil conditions and active plant–microbiome interactions may play a more dominant role in shaping plant traits than the historical legacy of microbial community composition alone (Burghardt et al. 2019; Liu et al. 2025).

Notably, we did not observe a single infected pea in the legacy experiment, despite inoculation with soil slurries from treatments where infection rates were highest. This absence of infection may reflect limitations of the slurry preparation method (e.g., loss of pathogen during settling), but more plausibly, it emphasizes the importance of nutrients and contaminants in “priming” infection. Unlike the direct exposure experiment, peas in the legacy experiment were nitrogen-limited and exposed to only trace levels of residual contaminants -- two factors that may have influenced their susceptibility. Additionally, due to limited slurry volumes, peas were only inoculated after germination, precluding our ability to detect microbiome effects on early developmental stages. Together, these factors support the hypothesis that pea infection is contingent not only on microbial presence but also on environmental context and timing of exposure.

That said, we did observe subtle legacy effects, particularly in microbiomes previously exposed to high contaminant concentrations, which were associated with reduced shoot:root ratios— a classic indicator of plant stress often linked to nutrient limitation or pathogen pressure (Agathokleous et al. 2019; Lopez et al. 2023; Seidel et al. 2024). Such patterns suggest that contaminant-induced microbiome disruption may contribute to pea stress responses, potentially through impaired nutrient cycling or altered microbial signaling (Liu et al. 2020). While these subtle effects were modest, they could have longer-term implications for plant health and resilience, particularly in systems where crops are repeatedly exposed to legacy-impacted soils. These findings open the door to future experiments that explicitly test the functional consequences of microbiome legacy. For example, a fully factorial design comparing plant responses to preconditioned versus naïve microbiomes, in the presence or absence of current soil treatments, could reveal whether legacy microbiomes buffer or exacerbate stress, offering insights into the potential for microbiome-mediated resilience in rotational cropping systems.

Our results also raise questions about microbiome resilience: to what extent do microbial communities recover from amendment- or contaminant-induced disturbances, and how quickly do they return to a functional state that supports plant health? In systems where biosolids or reclaimed water are applied intermittently, understanding the persistence or reversibility of microbiome shifts will be key to predicting long-term impacts on crop productivity. Ultimately, this work highlights the importance of considering both current and historical soil conditions when evaluating the risks and benefits of alternative amendments. While legacy effects may not always translate into immediate performance declines, their influence on early-stage traits and microbiome dynamics could shape plant responses over successive growing seasons. Future studies should explore how these legacy effects interact with crop rotation, soil type, and microbial recovery processes to inform sustainable amendment practices.

## Conclusion

Our study highlights the complex and crop-specific ways in which alternative soil amendments and their associated contaminants influence plant–microbiome interactions. By comparing multiple crop species under identical soil treatments, we show that both microbiome responses and plant trait outcomes vary markedly depending on crop identity. Notably, peas, reliant on symbiotic nitrogen fixation, exhibited heightened sensitivity to biosolid amendments—showing pronounced microbiome disruption and stress-related trait shifts. These fundings underscore the importance of functional plant traits in mediating amendment impacts.

While soil treatments induced clear and sometimes persistent changes in microbiome diversity and composition, these shifts did not uniformly translate into altered plant performance. Microbiome diversity, particularly in peas, was initially reduced but recovered over time, suggesting a degree of microbial resilience that may buffer long-term impacts. This recovery likely contributed to the limited effects observed in our legacy experiment, where microbiomes shaped by prior exposure had minimal influence on pea growth in the absence of ongoing stressors. Additionally, the timing of inoculation—post-germination—may have precluded early developmental impacts, further emphasizing the importance of exposure timing in microbiome–plant interactions.

Together, these findings demonstrate that amendment-induced microbiome shifts do not translate into uniform plant outcomes. Instead, they point to complex, context-dependent interplay shaped by crop identity, amendment type, contaminant load, and timing of exposure. As alternative amendments such as biosolids and reclaimed water become more common, our results highlight the need to move beyond one-size-fits-all assessments. Evaluating amendment impacts through the lens of crop-specific functional traits—such as symbiotic capacity and nutrient acquisition strategies—will be essential for predicting plant responses and designing resilient, sustainable agroecosystems.

## Acknowledgements

This work was supported by NSERC Discovery (RTD: RGPIN-2023-05144; GS: 2019-06456), the Canadian Foundation for Innovation (RTD: CFI42856), and McMaster University (RTD). We thank Zahava Dworkin for experimental assistance, Drs. Brian Golding, Ian Dworkin, and Karen Kidd for their guidance on data analysis and interpretation, and to Dr. Susan Dudley and the McMaster Data Lunch group for statistical advice.

## Author Contributions

Following CRediT System ([link](https://www.elsevier.com/researcher/author/policies-and-guidelines/credit-author-statement)) - Conceptualization: II, HS, RTD, and GS. Methodology: II, HS, and CC. Validation: II, HS, RTD, and GS. Formal analysis: II, RTD, and EC. Investigation: II, HS, and EC. Resources: RTD and GS. Data curation: II, RTD, and EC. Writing: II and RTD. Review and editing: HS, GS, EC, and CC. Visualization: II, RTD and EC. Supervision: RTD and GS.

## Data Availability

All raw sequencing reads are being deposited in NCBI’s Sequence Read Archive (SRA) under a BioProject accession number that will be provided prior to submission. All Supplementary Files will be uploaded to Dryad prior to submission for peer review. All experimental data and analysis code will be made publicly available on GitHub at the time of publication or can be provided upon request during peer review.

## Supplementary Information

Additional supporting information may be found in the online version of this article.

## Supplementary Files

**File S1: Contrasts from linear models testing main and interactive effects of crop species and amendment type (without additional contaminants), and contaminant concentration (spiking level, SL) on plant traits in the direct exposure experiment.**

**File S2: Contrasts from linear mixed models testing main and interactive effects of crop species, amendment type or contaminant concentration (SL 0-2), and timepoint on microbiome diversity.**

**File S3: Full model results from dbRDA (CAPSCALE) models testing main and interactive effects of crop species, soil treatment (amendment + contaminant concentration), timepoint, on microbiome composition. Arrows indicate plant traits significantly associated 981 with microbiome diversity.**

## Supplementary Tables

**Table S1:**
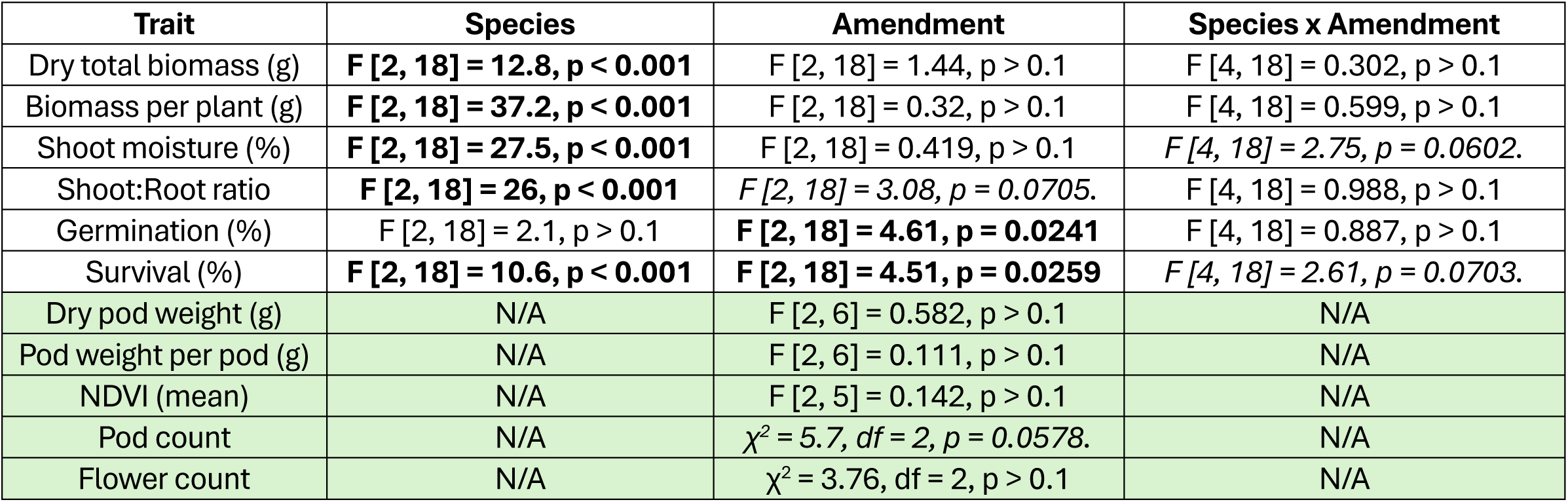
ANOVA results for linear models testing the effects of amendment type, crop species, and their interaction on plant traits measured in direct exposure experiment. Plant traits were measured across three crop species (lettuce, pea, radish) growing under three soil amendment treatments (biosolids, reclaimed water, and unamended controls). No additional contaminants added. F-statistics and p-values for main effects of species and amendment, as well as their interaction are reported. Cells shaded in green are for pea-specific traits. Significant (p < 0.05) terms are shown in bold, marginally significant (p < 0.1) terms in italics. Significant (p < 0.05) and marginally significant (0.05 < p < 0.1) contrasts (treatment vs control) are included where applicable, with effect sizes and confidence intervals. Significant (p < 0.1) contrasts are reported in Supplementary File S1.

**Table S2:** ANOVA results for linear models testing the effects of crop species, contaminant concentration (spiking level, SL) and their interaction on plant traits measured in direct exposure experiment. Plant traits were measured across three crop species (lettuce, pea, radish) grown under two amendment types (BS = biosolids; RW = reclaimed water) and exposed to three contaminant spiking levels (0, 1, 2). Unamended controls excluded from models. F-statistics and p-values for main effects of species and amendment, as well as their interaction are reported. Cells shaded in green are for pea-specific traits. Significant (p < 0.05) terms shown in bold, marginally significant (p < 0.1) terms in italics. Significant (p < 0.1) contrasts are reported in **Supplementary File S1**.

| Trait | Amend-<br>ment | Species | Spiking level (SL) | Species x SL |
| --- | --- | --- | --- | --- |
| Dry total biomass (g) | RW | **F [2, 18] = 41.9, $p < 0.001$** | F [2, 18] = 0.505, $p > 0.1$ | *F [4, 18] = 2.37, $p = 0.0908$* |
| | BS | **F [2, 17] = 32.2, $p < 0.001$** | F [2, 17] = 0.256, $p > 0.1$ | F [4, 17] = 1.78, $p > 0.1$ |
| Biomass per plant (g) | RW | **F [2, 18] = 69.8, $p < 0.001$** | F [2, 18] = 1.49, $p > 0.1$ | **F [4, 18] = 4.2, $p = 0.0142$** |
|  | BS | **F [2, 17] = 22, $p < 0.001$** | **F [2, 17] = 4.69, $p = 0.0239$** | *F [4, 17] = 2.77, $p = 0.0608$* |
| Shoot moisture (%) | RW | **F [2, 18] = 34.2, $p < 0.001$** | F [2, 18] = 0.0038, $p > 0.1$ | **F [4, 18] = 4.22, $p = 0.0139$** |
| | BS | **F [2, 17] = 67.4, $p < 0.001$** | F [2, 17] = 1.94, $p > 0.1$ | F [4, 17] = 0.406, $p > 0.1$ |
| Shoot:Root ratio | RW | **F [2, 18] = 32.3, $p < 0.001$** | F [2, 18] = 0.642, $p > 0.1$ | F [4, 18] = 0.943, $p > 0.1$ |
| | BS | **F [2, 17] = 18.8, $p < 0.001$** | F [2, 17] = 0.793, $p > 0.1$ | F [4, 17] = 0.755, $p > 0.1$ |
| Germination (%) | RW | F [2, 18] = 2.58, $p > 0.1$ | **F [2, 18] = 5.23, $p = 0.0162$** | F [4, 18] = 0.316, $p > 0.1$ |
| | BS | **F [2, 18] = 38.1, $p < 0.001$** | F [2, 18] = 1.98, $p > 0.1$ | **F [4, 18] = 4.3, $p = 0.0129$** |
| Survival (%) | RW | **F [2, 18] = 8.2, $p = 0.00294$** | **F [2, 18] = 3.89, $p = 0.0393$** | F [4, 18] = 0.228, $p > 0.1$ |
|  | BS | **F [2, 18] = 60.9, $p < 0.001$** | *F [2, 18] = 3.36, $p = 0.0576$* | **F [4, 18] = 3.08, $p = 0.0425$** |
| Dry pod weight (g) | RW | N/A | F [2, 6] = 1.26, $p > 0.1$ | N/A |
| | BS | N/A | F [2, 4] = 0.0228, $p > 0.1$ | N/A |
| Pod weight per pod (g) | RW | N/A | F [2, 6] = 0.7, $p > 0.1$ | N/A |
| | BS | N/A | F [2, 4] = 2.16, $p > 0.1$ | N/A |
| Pod count | RW | N/A | **$\chi^2 = 6.31$, $df = 2$, $p = 0.0426$** | N/A |
|  | BS | N/A | **$\chi^2 = 14.9$, $df = 2$, $p < 0.001$** | N/A |
| Flower count | RW | N/A | $\chi^2 = 0.0496$, $df = 2$, $p > 0.1$ | N/A |
| | BS | N/A | $\chi^2 = 1.3$, $df = 2$, $p > 0.1$ | N/A |

**Table S3:** ANOVA results for linear mixed models testing the effects of amendment type, crop species, timepoint, and their interactions on microbiome diversity measured in direct exposure experiment. Shannon and Inverse Simpson diversity indices were calculated for microbiomes sampled at two timepoints (1 wk PP: one-week post-planting; Harvest), across three crop species (lettuce, pea, radish) growing under three soil amendment treatments (biosolids, reclaimed water, and unamended controls). No additional contaminants added. Chi-squared statistics and p-values for main and interactive effects are reported. Significant (p < 0.05) terms are shown in bold, marginally significant terms (p < 0.1) in italics. Significant (p < 0.1) contrasts reported in **Supplementary File S2**.

| Term | Shannon | Inverse Simpson |
| --- | --- | --- |
| Species | $\chi^2 [2] = 5.44, p = 0.0659$ | $\chi^2 [2] = 1.51, p > 0.1$ |
| Amendment | $\chi^2 [2] = 0.461, p > 0.1$ | $\chi^2 [2] = 1.35, p > 0.1$ |
| Timepoint | **$\chi^2 [1] = 12.2, p < 0.001$** | **$\chi^2 [1] = 23.2, p < 0.001$** |
| Species x Amendment | $\chi^2 [4] = 7.09, p > 0.1$ | $\chi^2 [4] = 3.15, p > 0.1$ |
| Species x Timepoint | **$\chi^2 [2] = 6.84, p = 0.0327$** | $\chi^2 [2] = 4.73, p = 0.0941$ |
| Amendment x Timepoint | $\chi^2 [2] = 0.364, p > 0.1$ | $\chi^2 [2] = 3.86, p > 0.1$ |
| Species x Amendment x Timepoint | **$\chi^2 [4] = 14.2, p = 0.00658$** | $\chi^2 [4] = 6.45, p > 0.1$ |

**Table S4:** ANOVA results for linear mixed models testing the effects of crop species, contaminant concentration, timepoint, and their interactions on microbiome diversity measured in direct exposure experiment. Shannon and Inverse Simpson diversity indices were calculated for microbiomes sampled at two timepoints (1 wk PP: one-week post-planting; Harvest), across three crop species (lettuce, pea, radish) grown under two amendment types (BS = biosolids; RW = reclaimed water), each being exposed to three contaminant spiking levels (0, 1, 2). Unamended controls excluded from models. Significant (p < 0.05) terms are shown bold, marginally significant terms (p < 0.1) in italics. Significant (p < 0.1) contrasts reported in **Supplementary File S2**.

| Term | Shannon (RW) | Shannon (BS) | Inverse Simpson (RW) | Inverse Simpson (BS) |
| --- | --- | --- | --- | --- |
| Species | $\chi^2 [2] = 1.77, p > 0.1$ | **$\chi^2 [2] = 11.7, p = 0.00283$** | $\chi^2 [2] = 1.15, p > 0.1$ | **$\chi^2 [2] = 8.21, p = 0.0165$** |
| Spiking level (SL) | $\chi^2 [2] = 0.923, p > 0.1$ | $\chi^2 [2] = 0.511, p > 0.1$ | $\chi^2 [2] = 1.01, p > 0.1$ | $\chi^2 [2] = 0.784, p > 0.1$ |
| Timepoint | **$\chi^2 [1] = 19.2, p < 0.001$** | **$\chi^2 [1] = 6.84, p = 0.00889$** | **$\chi^2 [1] = 45.8, p < 0.001$** | **$\chi^2 [1] = 27.8, p < 0.001$** |
| Species x SL | $\chi^2 [4] = 9.26, p = 0.0549$ | $\chi^2 [4] = 1.29, p > 0.1$ | $\chi^2 [4] = 6.6, p > 0.1$ | $\chi^2 [4] = 1.37, p > 0.1$ |
| Species x Timepoint | $\chi^2 [2] = 2.08, p > 0.1$ | **$\chi^2 [2] = 6.59, p = 0.0371$** | $\chi^2 [2] = 1.91, p > 0.1$ | $\chi^2 [2] = 5.2, p = 0.0741$ |
| SL x Timepoint | $\chi^2 [2] = 0.547, p > 0.1$ | $\chi^2 [2] = 2.47, p > 0.1$ | $\chi^2 [2] = 1.18, p > 0.1$ | $\chi^2 [2] = 1.84, p > 0.1$ |
| Species x SL x Timepoint | $\chi^2 [4] = 4.48, p > 0.1$ | $\chi^2 [4] = 1.53, p > 0.1$ | $\chi^2 [4] = 2.12, p > 0.1$ | $\chi^2 [4] = 1.03, p > 0.1$ |

**Table S5:**
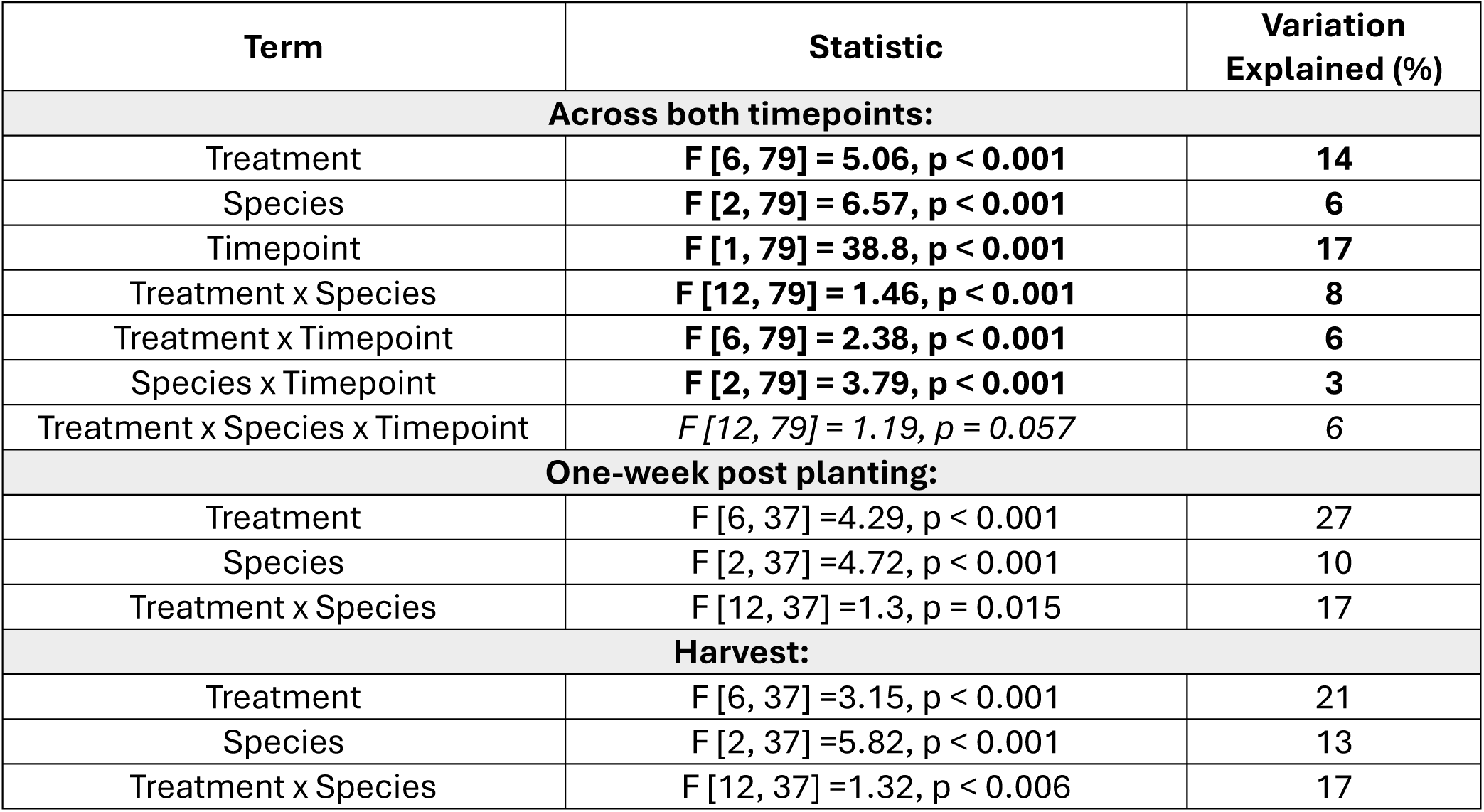
ANOVA-like results for dbRDA (CAPSCALE) analyses testing the effects of crop species, soil treatment (amendment + contaminant level), timepoint, and their interactions on microbiome composition in the direct exposure experiment. Bray-Curtis distances were calculated for microbiomes sampled at two timepoints (1 wk PP: one-week post-planting; Harvest), across three crop species (lettuce, pea, radish) grown under two amendment types (BS = biosolids; RW = reclaimed water) and three contaminant spiking levels (0, 1, 2), resulting in six treatments plus an unamended control. Significant (p < 0.05) terms are shown in bold, marginally significant terms (p < 0.1) in italics. Full model results available in **Supplementary File S3**.

| Term | Statistic | Variation Explained (%) |
| --- | --- | --- |
| **Across both timepoints:** |  |  |
| Treatment | **F [6, 79] = 5.06, $p < 0.001$** | **14** |
| Species | **F [2, 79] = 6.57, $p < 0.001$** | **6** |
| Timepoint | **F [1, 79] = 38.8, $p < 0.001$** | **17** |
| Treatment x Species | **F [12, 79] = 1.46, $p < 0.001$** | **8** |
| Treatment x Timepoint | **F [6, 79] = 2.38, $p < 0.001$** | **6** |
| Species x Timepoint | **F [2, 79] = 3.79, $p < 0.001$** | **3** |
| Treatment x Species x Timepoint | *F [12, 79] = 1.19, $p = 0.057$* | 6 |
| **One-week post planting:** |  |  |
| Treatment | F [6, 37] = 4.29, $p < 0.001$ | 27 |
| Species | F [2, 37] = 4.72, $p < 0.001$ | 10 |
| Treatment x Species | F [12, 37] = 1.3, $p = 0.015$ | 17 |
| **Harvest:** |  |  |
| Treatment | F [6, 37] = 3.15, $p < 0.001$ | 21 |
| Species | F [2, 37] = 5.82, $p < 0.001$ | 13 |
| Treatment x Species | F [12, 37] = 1.32, $p < 0.006$ | 17 |

## Supplementary Figures

**Figure S1:**
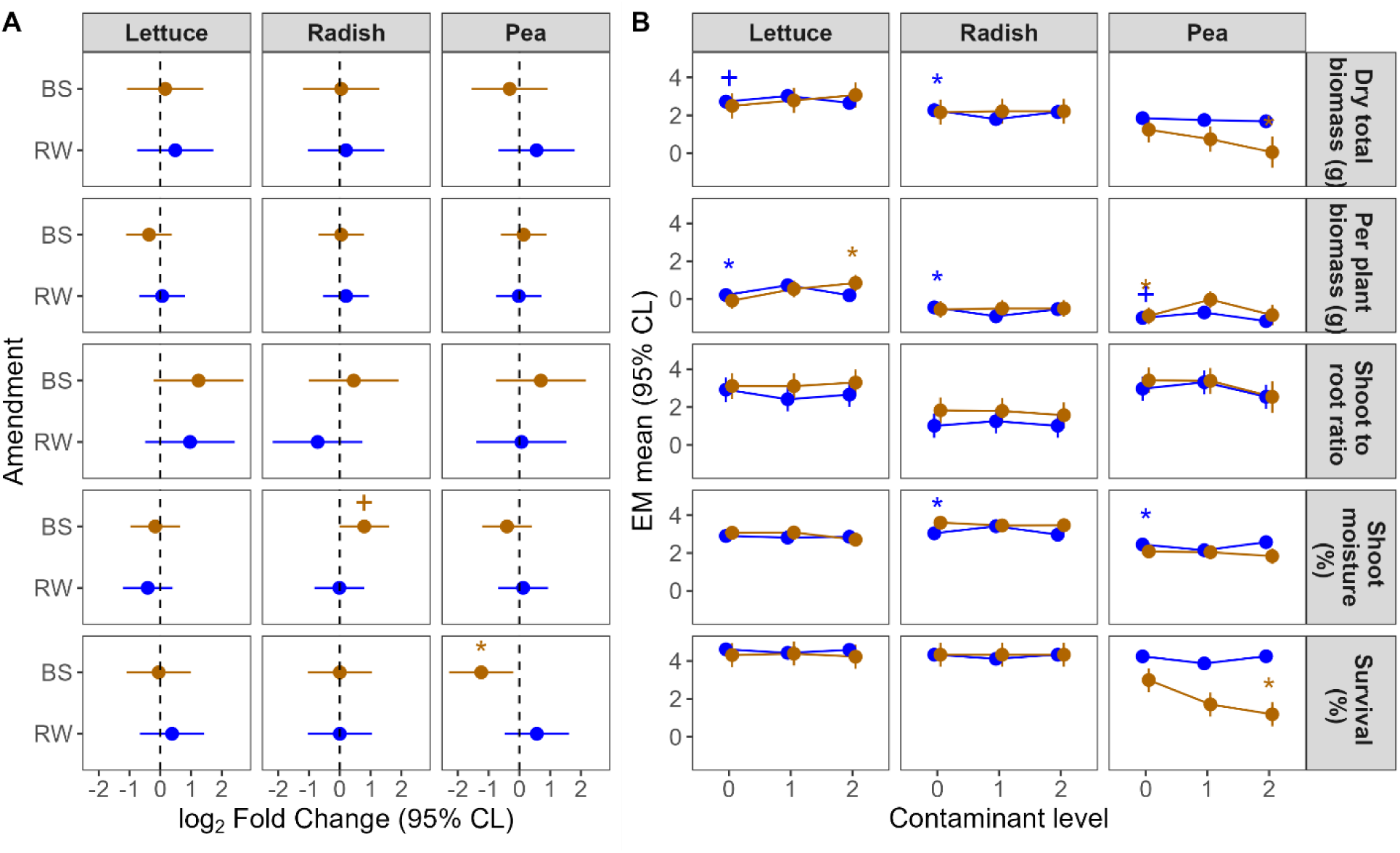
Pea germination and survival are negatively impacted by biosolid amendments with increasing contaminant concentrations. **A** Dunnet-style contrasts based on linear models (LMs) testing for the main and interactive effects of crop species and amendment type in the absence of added contaminants (spiking level 0; model set 1). Effect sizes (points) represent the log_2_ fold change (FC) of either biosolids (BS, in brown) or reclaimed water (RW, in blue) compared to unamended controls (dashed vertical lines). Error bars represent 95% confidence limits (CL). Asterisks (*) and plus signs (+) represent significant (p < 0.05) and marginally significant (0.05 < p < 0.1) contrasts, respectively. **B** Linear and quadratic estimates based on LMs testing for the main and interactive effects of crop species and contaminant spiking levels (0-2) for each amendment type separately (blue = RW; brown = BS). Estimated marginal means (points) are shown along with their corresponding 95% CL (error bars). Asterisks (*) or plus signs (+) represent significant (p < 0.05) or marginally significant (0.05 < p < 0.1), respectively, linear (right) or quadratic (left) estimates.

**Figure S2:**
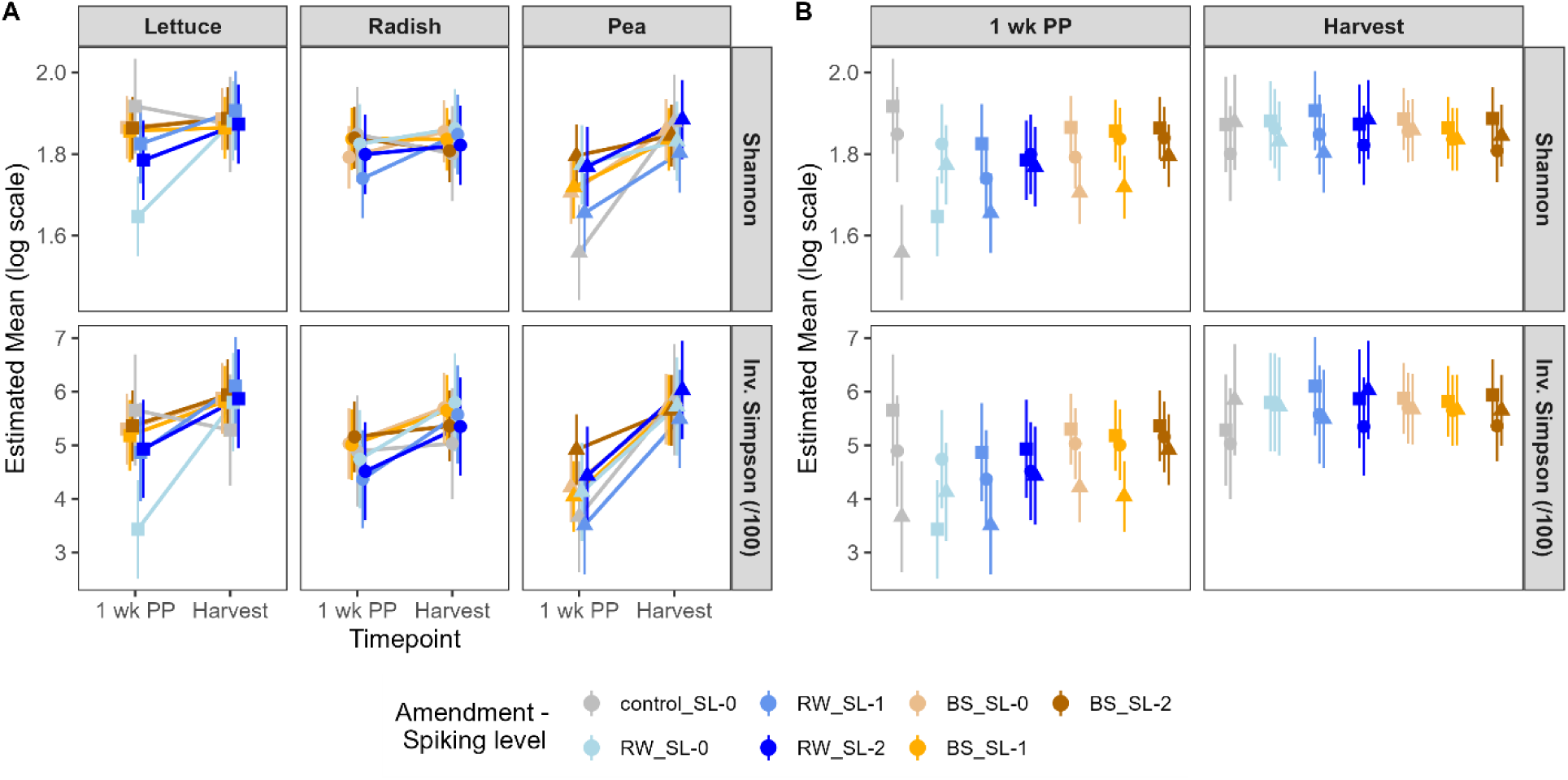
Species-driven differences in microbiome diversity diminish over time. **A,** Estimated marginal (EM) means of microbiome diversity for each crop species within each soil treatment across timepoints; error bars represent 95% confidence intervals. **B,** Same means redrawn to highlight species comparisons within each timepoint and treatment; error bars represent 95% confidence intervals.

**Figure S3:**
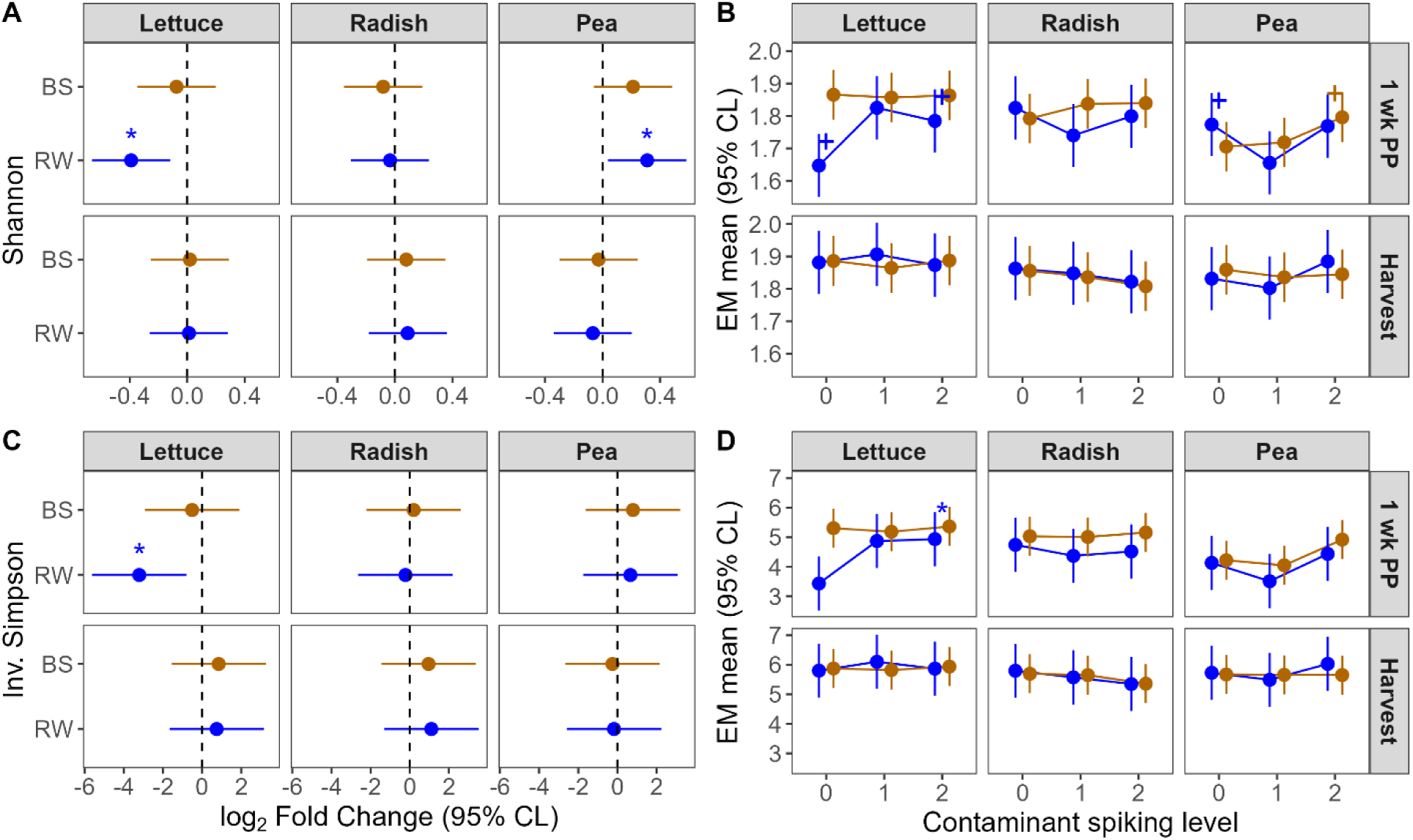
Microbiome diversity is impacted by amendment and contaminant concentrations in a crop-species specific manner. **A, C** Dunnet-style contrasts based on linear mixed models (LMMs) testing for the main and interactive effects of crop species, amendment type, and timepoint in the absence of added contaminants (spiking level 0) on Shannon (**A**) and Inverse Simpson (**C**) diversity metrics, each measured at two timepoints, one-week post-planting (top) and at harvest (bottom). Effect sizes (points) represent the log_2_ fold change (FC) of either biosolids (BS, in brown) or reclaimed water (RW, in blue) compared to unamended controls (dashed vertical lines). Error bars represent 95% confidence limits (CL). Asterisks (*) and plus signs (+) represent significant (p < 0.05) and marginally significant (0.05 < p < 0.1) contrasts, respectively. **B, D** Linear and non-linear (i.e., quadratic) responses to increasing contaminant concentrations based on LMMs testing for the main and interactive effects of crop species, contaminant spiking levels (0-2), and timepoint on Shannon (**B**) and Inverse Simpson (**D**) indices, each measured at two timepoints, one-week post-planting (top) and at harvest (bottom), for each amendment type separately (blue = RW; brown = BS). Estimated marginal means (points) are shown along with their corresponding 95% CL (error bars). Asterisks (*) or plus signs (+) represent significant (p < 0.05) or marginally significant (0.05 < p < 0.1), respectively, linear (right) or quadratic (left) estimates.

**Figure S4:**
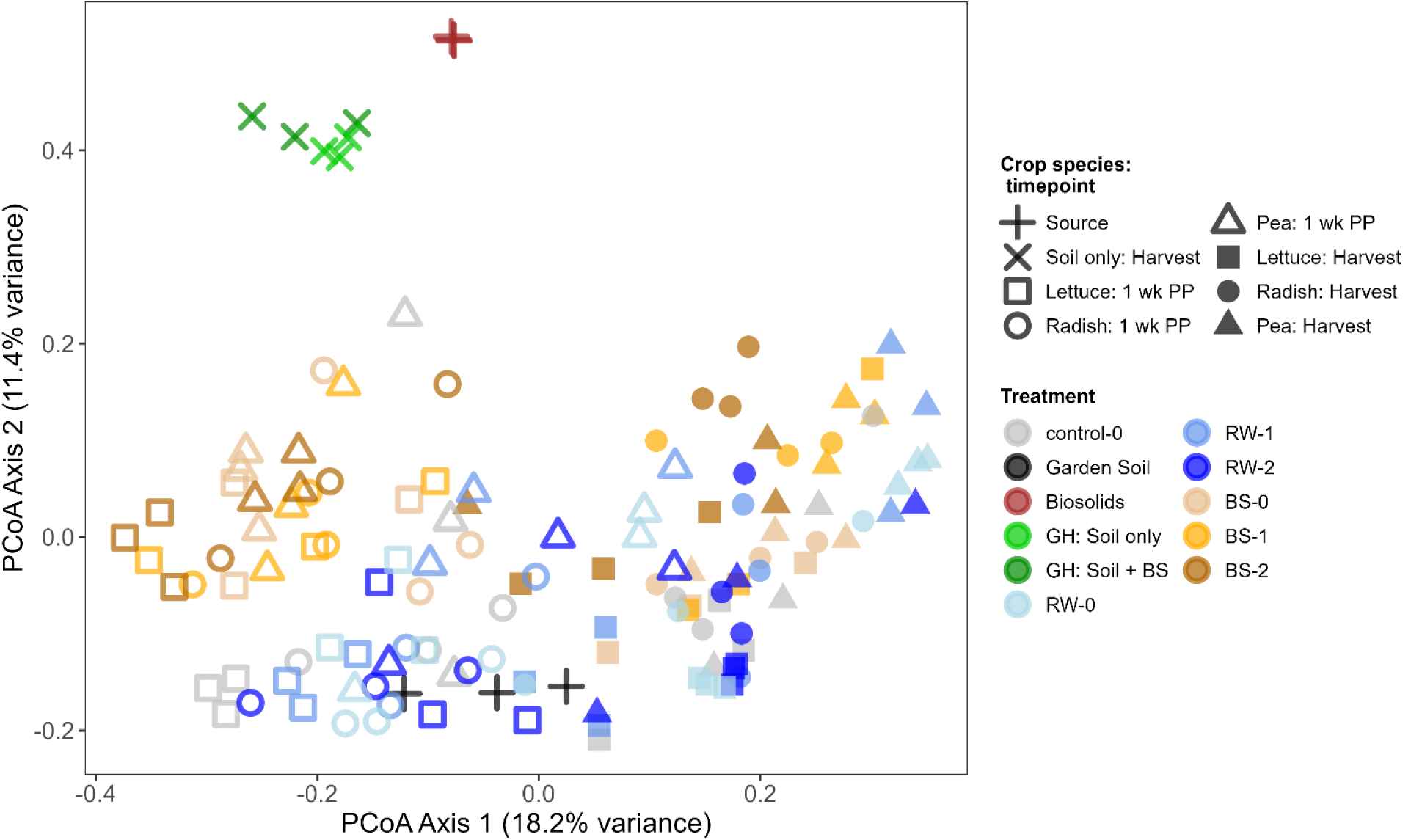
Microbiome composition of source versus experimental samples. **A,** Estimated marginal (EM) means of microbiome diversity for each crop species within each soil treatment across timepoints; error bars represent 95% confidence intervals. **B,** Same means redrawn to highlight species comparisons within each timepoint and treatment; error bars represent 95% confidence intervals.

**Figure S5:**
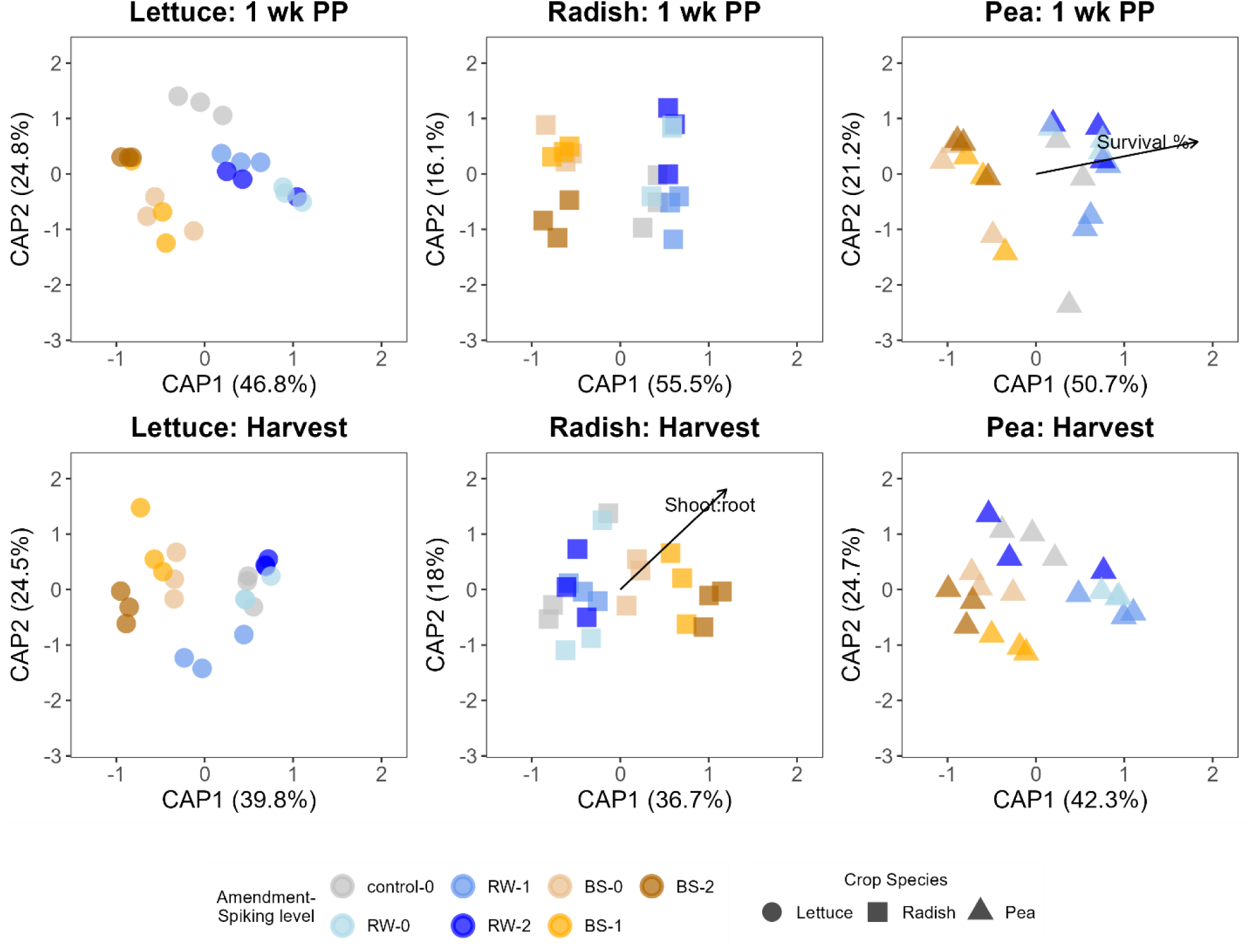
Distance-based redundancy analysis (dbRDA) ordination of Bray–Curtis dissimilarities constrained by plant traits, crop species, and soil treatments. Points represent individual samples coloured by amendment–spiking level treatment and shaped by crop species (Lettuce, Radish, Pea). Axes (CAP1 and CAP2) show the proportion of constrained variation explained by the first two canonical axes. Significant plant traits (p < 0.05, permutation test) are fitted as vectors, with arrow length proportional to correlation strength and direction indicating association with community composition. Forward selection was applied to identify the most parsimonious set of predictors.

## Supplementary Methods

**Methods S1:**
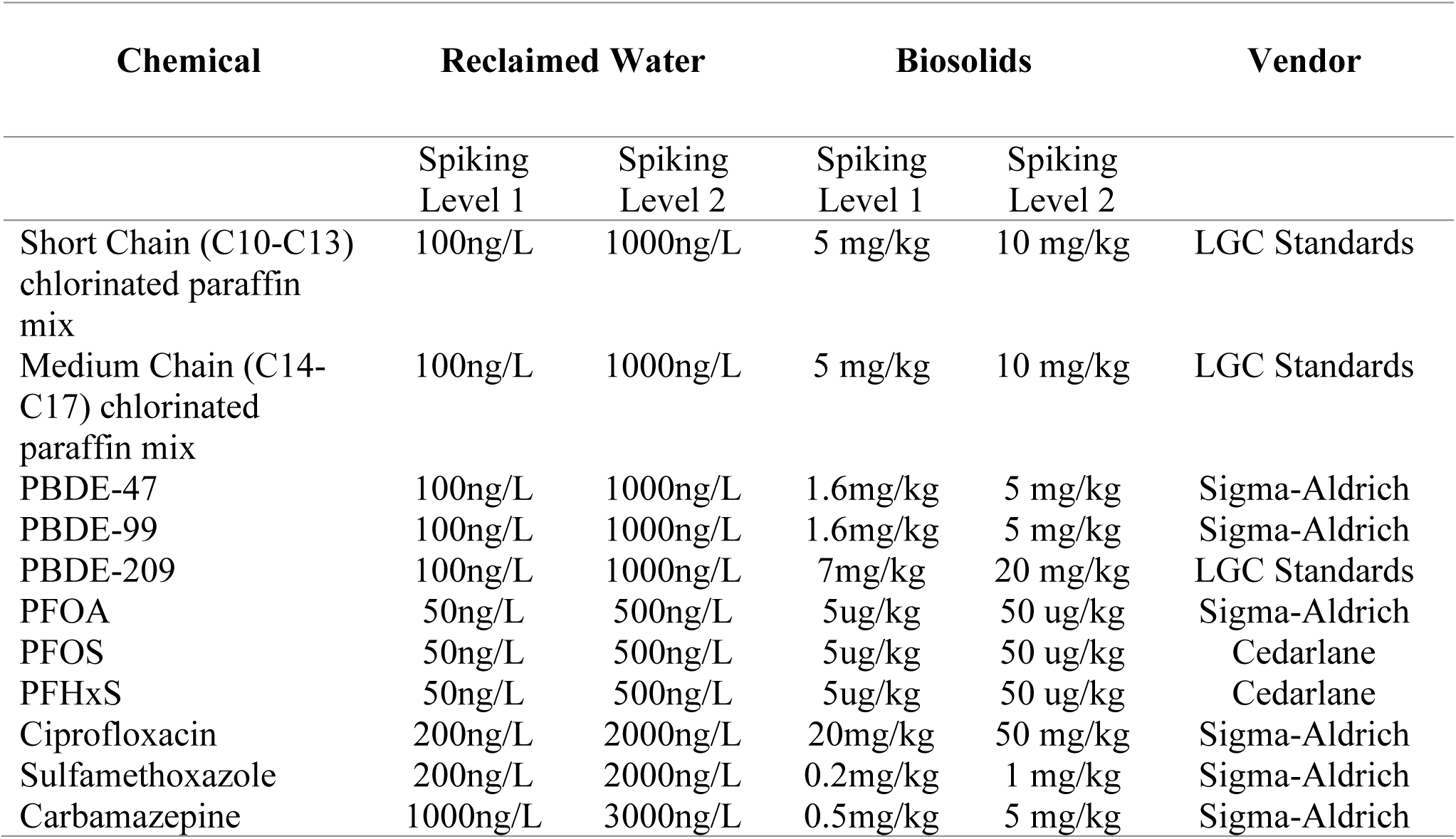
Contaminant mixtures.

**Methods S2:**
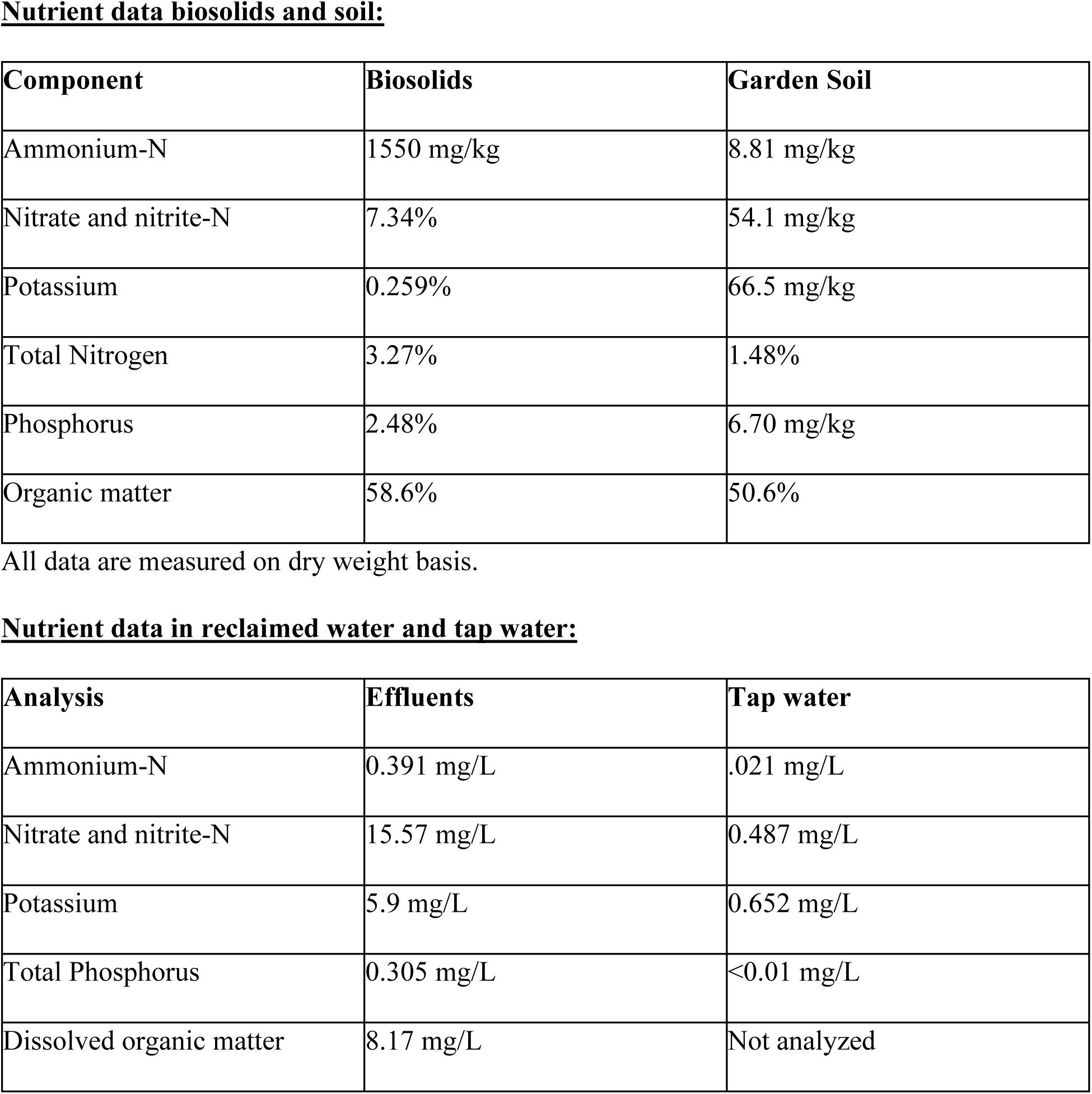
Nutrients present in source materials.

**Methods S3:**
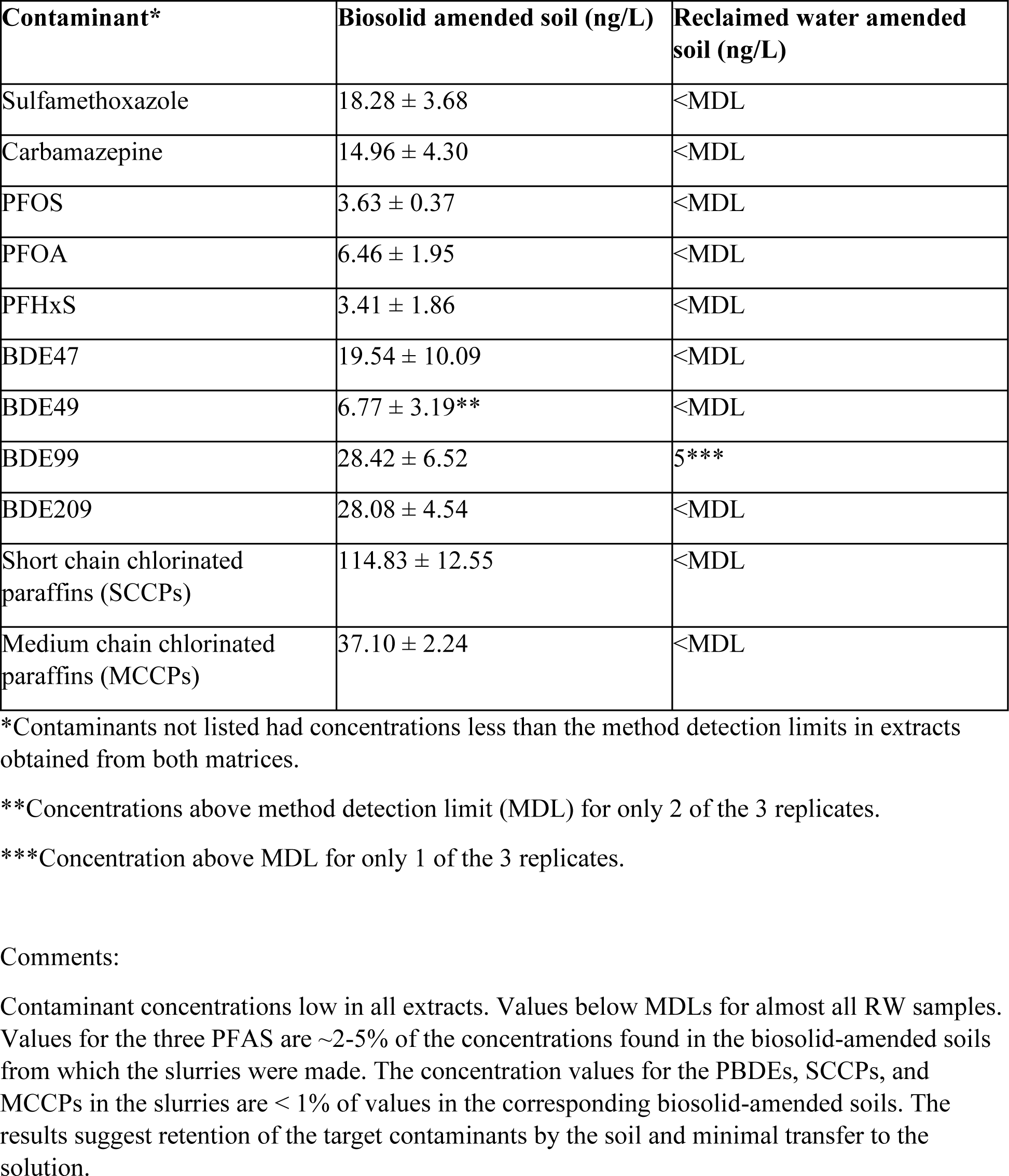
Contaminant levels in soil slurries.

## Notes

### Competing Interest Statement

The authors have declared no competing interest.

